# A spatiotemporal perturbational atlas of the post-pneumonectomy lung

**DOI:** 10.64898/2026.11.25.768918

**Authors:** Tomohiro Obata, Diego Perez Aracena, Maria Tomas Gracia, Sophie E. Edelstein, Satoshi Mizoguchi, Nuoya Wang, Natan-El A. Mensa Sowah, Colten Danelski, Connor Haynes, Olivia Porcaro, Naoya Kitamura, Katarina Lile, William Dorst, Allison M. Greaney, Taylor Adams, Rachel Rivero, Hahram Kim, Vi Lee, Junchen Yang, Yifan Yuan, Jonas C. Schupp, Andre Levchenko, Aaron Osgood-Zimmerman, David H. Stitelman, Tomoshi Tsuchiya, Themis Kyriakides, Naftali Kaminski, Laura E. Niklason, Micha Sam Brickman Raredon

## Abstract

Perturbational spatial atlases have begun to map regenerative dynamics in a variety of tissues, yet lung remodeling has not been characterized at comparable spatial and temporal resolution. Post-pneumonectomy lung regeneration (PPLR) provides a tractable mammalian model for tissue regeneration that can be studied in situ. Here, we present a spatiotemporal perturbational atlas of PPLR in *Rattus norvegicus*, combining single-cell RNA sequencing (scRNAseq), high-resolution spatial transcriptomics (10x Genomics Xenium), and NICHES-based spatial tissue connectomics. We observe three distinct tissue architectures at successive timepoints after pneumonectomy, each characterized by coordinated shifts in local cellular composition and intercellular signaling. At Day 3 (D3), a line of interstitial macrophages colocalizes directly beneath an intact mesothelium among active mechanotransductive and inflammatory signaling. We also observe the unexpected appearance of a transitional remodeling cellular community at D7 which appears largely absent at later timepoints, marked by fibrosis-mimetic signaling. At D14, an alveologenic architecture at distal lung tissue displays a connectivity signature consistent with active de novo growth, distinct from baseline homeostasis. Our findings establish a spatiotemporal map of multicellular architecture during regenerative lung remodeling and introduce a generalizable framework for spatial connectomic analysis to be used in perturbational tissue systems biology.

## Introduction

Perturbational strategies in systems biology study how biological systems respond when deliberately pushed away from a set of baseline homeostatic conditions to uncover cause-and-effect relationships(Ideker et al., 2001; Kotas and Medzhitov, 2015). The field now spans genetic-perturbation using at cellular (Dixit et al., 2016; Replogle et al., 2022), spatial (Dhainaut et al., 2022; Shen et al., 2026), and whole-tissue in vivo (Saunders et al., 2025) modalities, alongside computational efforts on perturbation-response prediction (Bunne et al., 2024; Bunne et al., 2023; Lotfollahi et al., 2023); in parallel, gross physiological and surgical perturbations have been measured as creating whole organ and tissue responses that can also be interrogated through multi-modal investigation (Li et al., 2025; Polonsky et al., 2024; Wu et al., 2024). Tissue homeostasis, stress responses, and inflammation form a continuum of responses to such perturbations (Chovatiya and Medzhitov, 2014; Meizlish et al., 2021). The positions on this continuum within a tissue at a given moment depends on the magnitude of perturbation away from tightly controlled homeostatic setpoints and on whether homeostatic effectors can adapt to the perturbation (Goldstein and Kopin, 2007). Perturbational tissue spatial atlases offer a way to read these positions directly *in situ* (Rood et al., 2024).

Such atlases have already been built for several organs responding to defined perturbation sets. In skin, a 4D multimodal atlas of mammalian wound healing has reconstructed repair phase transitions using integrated single-cell and high-definition spatial transcriptomics (Cheong et al., 2026). In liver, a spatiotemporal atlas of partial-hepatectomy regeneration has mapped molecular gradients and intercellular signaling networks coordinating regeneration (Xu et al., 2024). These atlases involve tissue-, organ-, and organism-scale perturbations that differ from traditional cell-state atlases and targeted genetic studies along traditional developmental trajectories. They both map non-traditional morphogenesis as multicellular communities assemble and dissolve in space and time, while simultaneously helping to identify the specific regulated variables that holistically guide tissue architecture. These regulated variables are maintained by minimal tissue units, defined as groups of cell populations whose coordinated activities sustain tissue homeostasis. Only a small number of such variables, including oxygen tension and pH, are currently understood well enough to link changes in their regulation to tissue-scale responses such as angiogenesis and acid-base control (Adler et al., 2023; Meizlish *et al*., 2021; Yonassi et al., 2025). Some key variables known to be involved in guiding the lung tissue’s response post-pneumonectomy are tissue tension and strain (Hsia, 2017; Mentzer, 2018; Ysasi et al., 2017).

A growing literature synthesizes the lung as a truly ‘re-generative’ organ. Compensatory growth has been documented after experimental pneumonectomy in mice (Peng et al., 2025; Voswinckel et al., 2004), rats (Addis, 1928; Brody, 1975; Cagle et al., 1990), rabbits (Das and Thurlbeck, 1979; Langston et al., 1977; Thurlbeck and Langston, 1989), ferrets (McBride, 1989), beagles (Hsia et al., 1994; Ohata et al., 2022), and pigs (Dao et al., 2018), and has been observed clinically in young humans following lobectomy (Shikuma et al., 2018; Wakamatsu et al., 2020). In rodents specifically, this process involves local tissue hyperplasia (Cagle *et al*., 1990), and for this reason, we refer to the remodeling process as a whole as Post-Pneumonectomy Lung Regeneration (PPLR).

PPLR begins within days of surgery and combines cell proliferation, reactivation of developmental signals that drive branching morphogenesis and alveologenesis (Brody, 1975; Brody et al., 1978; Cagle and Thurlbeck, 1988; Fisher and Simnett, 1973), and coordinated expansion of the airway and vascular tree (Kotton and Morrisey, 2014). It shares mechanistic features with regenerative programs in other organ systems, including initial exit from homeostasis, activation of developmental patterns, and tissue maturation (Kho et al., 2013; Moya and Halder, 2019; Mui et al., 2026; Tsissios et al., 2026). At the tissue scale, PPLR proceeds through both neoalveolarization and expansion of existing alveolar units (Fehrenbach et al., 2008; Rippa et al., 2021; Voswinckel *et al*., 2004). This response is beginning to be sufficiently well established that it is cited as a neo-alveologenesis model in other experiments (Konkimalla et al., 2023).

Previous work has established that PPLR is triggered in large part by mechanotransduction of elevated tissue strain following pneumonectomy (Gilbert and Rannels, 1998; Hsia, 2017), while neither transient hypoxia nor altered pulmonary blood flow is sufficient on its own to engender the remodeling response (Karl et al., 1988; McBride et al., 1992). Notably, this mechanotransduction converges on YAP activation in alveolar stem cells, which is required for their proliferation and differentiation during post-pneumonectomy regeneration (Liu et al., 2016); endothelial YAP1 is additionally required for compensatory vascular and alveolar morphogenesis (Mammoto and Mammoto, 2019). Critically, the same mechanical-tension axis can drive the opposite outcome: when alveolar stem cells cannot regenerate, sustained tension instead activates a profibrotic TGFβ program (Lechner et al., 2017), positioning tension as finely tuned global and local homeostatic variable for maintaining lung tissue morphology.

Beyond mechanotransduction, PPLR also requires a specific immune contribution; recruited CCR2+ monocytes and Type-2 immunity are known to support alveolar epithelial type-2 cell proliferation during regeneration (Lechner *et al*., 2017), and recent work suggests group 2 innate lymphoid cells are themselves mechanosensitive via Piezo1, raising the possibility that the mechanical and immune axes of PPLR are coupled (Lim et al., 2025). Despite this growing characterization of the multicellular involvement in PPLR, this process has not been profiled in a high-resolution spatiotemporal manner, leaving unmapped how the remaining lung grows to a new size to accommodate the relative strain increase imposed post-pneumonectomy.

Investigating tissue biology under forced gross perturbation such as pneumonectomy is inherently complex; many sensor-effector circuits are obfuscated by the highly multidimensional, multi-agent response, as the agential material (Davies and Levin, 2023) finds a way to resolve competing demands (Levin, 2023). Snapshot profiling captures discrete points along this continuous process, raising hypotheses about which homeostatic variables are being regulated without yet counterfactually establishing them. Additionally, the clean cell-type categories visible at homeostasis blur during regenerative remodeling, as cells across multiple lineages exit their canonical states and enter transitional configurations often reminiscent of their developmental programs (Beers and Morrisey, 2011; Ke et al., 2025a; Viragova et al., 2024). Single-cell studies of regenerative lung remodeling have already begun to characterize this phenomenon: a transitional epithelium and fibroblast pair has been described as co-localizing during lung repair (Konkimalla *et al*., 2023), and analogous transitional populations have been recovered across regenerative contexts (Ayyaz et al., 2019; Ke and Cardoso, 2026; Marconi et al., 2021; Sicklick et al., 2006). The spatial emergence and resolution of these transitional ensembles within a single regenerating lung, however, has not been directly mapped.

Distinguishing productive from pathological transitions through a diverse tissue state landscape requires reading not just which cells are present, but which signals connect and reinforce unique architectures. Spatial transcriptomics now allows multicellular communities to be profiled directly *in situ* (Janesick et al., 2023; Larsson et al., 2021; Rao et al., 2021), and connectomic methods such as NICHES (Raredon et al., 2023) extend this readout to ligand-receptor signaling at the level of individual cell-pair edges, treating the tissue as a graph of communicating cells rather than a catalog of isolated cell types. Applied to a perturbational spatial atlas, this inferred connectomic readout lets a static spatial snapshot be interpreted through the lens of known signaling mechanisms. Because the ligand-receptor literature describes not just isolated gene expression but mechanistic interactions, the connectomic view localizes these functionally characterized pathways within the cellular ensembles present at each timepoint.

In this study, we use imaging-based spatial transcriptomics and tissue connectomics to study PPLR for the first time. Our findings reveal three distinct parenchymal architectures at successive timepoints after pneumonectomy (D3, D7, and D14, denoting 3-, 7-, and 14-days post-surgery, relative to a D0 baseline), each defined by characteristic cellular ensembles and local cell–cell communication circuits, together consistent with phases of a coordinated remodeling process. Our atlas provides unprecedented insight into the multicellular character of tissue-scale remodeling during PPLR and offers a framework to ask how the same cellular and signaling primitives produce regeneration in one tissue context and pathology in another.

## Results

### 1. A spatiotemporal atlas resolves the post-pneumonectomy lung across four timepoints

Tissue was collected from residual lung tissue undergoing regeneration following pneumonectomy (Fig 1a, Fig 1b) and prepared for both scRNAseq (DropSeq, 10x genomics v2 and v3 chemistries) and Xenium imaging-based spatial transcriptomics (Fig1c). Whole lobes were perfusion-fixed (to allow for capillary visualization) and processed for H&E and Trichrome staining (Fig S1). Selected histologic findings from these experiments are shown in Fig 1d and summarized here. At D3, the lung parenchyma was observed to be broadly inflamed, displaying adventitial, subpleural, and parenchymal mononuclear infiltration, intra-alveolar septal thickening, and notable changes in the mesothelial and subpleural architecture. Spatially heterogeneous remodeling was then observed at D7, characterized by the formation of matrix-dense and microvasculature-sparse intra-parenchymal, subpleural, and peri-adventitial regions. By D14, signs of alveologenic ‘secondary septation’ (or ‘alveolar ringing’ (Warburton, 2021)) were observed broadly throughout the lung, characterized histologically by the appearance of nascent alveolar constriction rings extruding from the walls of alveolar ducts (Fig 1d and Fig S1).

**Figure 1.**
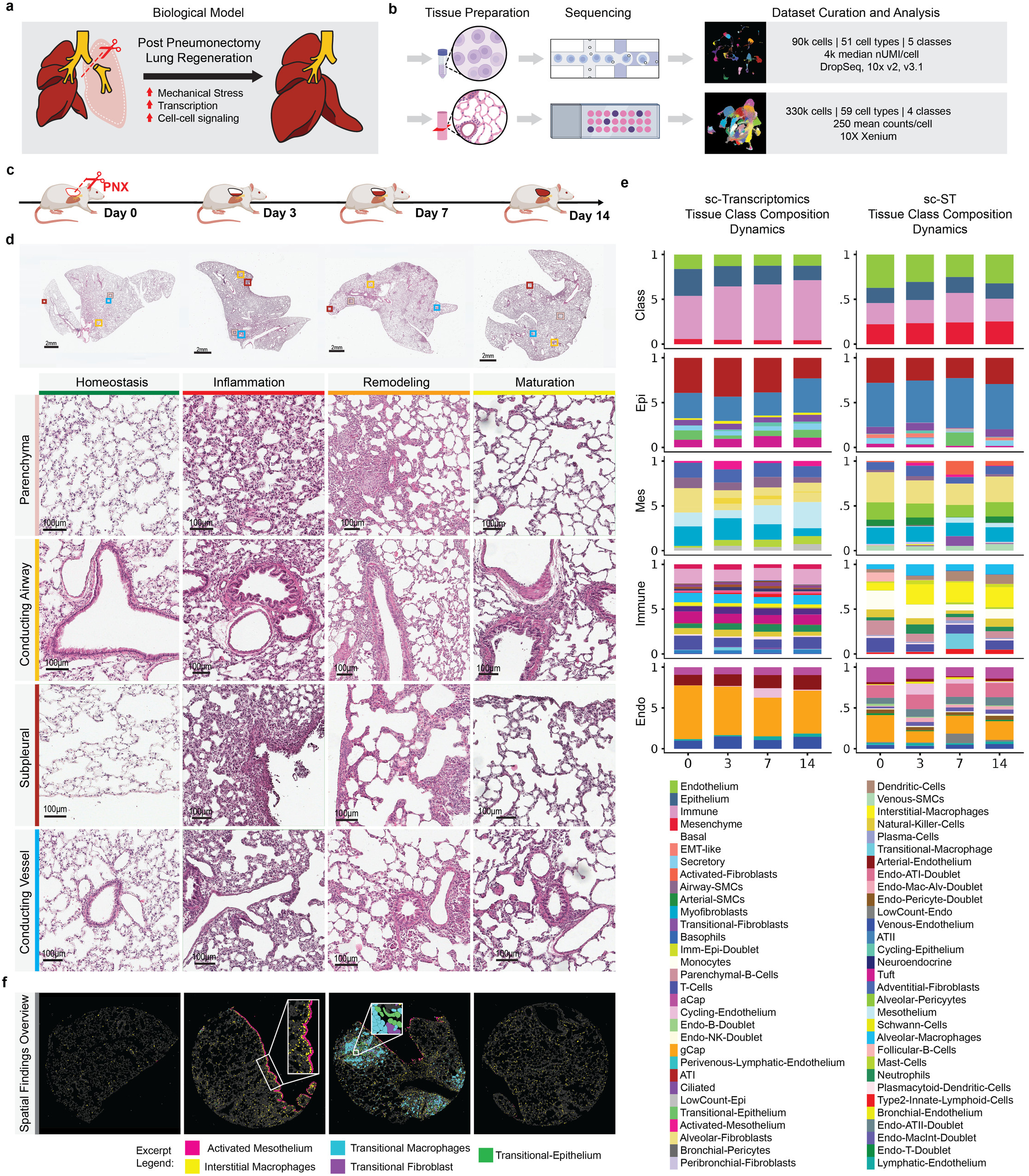
Paired single-cell and spatial transcriptomics resolve the cellular and tissue-level dynamics of post-pneumonectomy lung regeneration in *Rattus Norvegicus*. **(A) Biological model.** Pneumonectomy of the left lung in adult *Rattus Norvegicus* drives compensatory growth of the residual lobes. The process is accompanied by increased mechanical stress, transcriptional activation, and intercellular signaling. **(B) Tissue processing, sequencing, and dataset curation.** The post-pneumonectomy accessory lobe was profiled by two complementary modalities. Top: droplet-based single-cell RNA sequencing (DropSeq, 10x Chromium v2 and v3.1) yielded ∼90,000 cells annotated into 51 cell types across 5 lineage classes (∼4,000 median nUMI/cell). Bottom: 10x Genomics Xenium imaging-based spatial transcriptomics on a 488-gene custom panel yielded ∼330,000 cells annotated into 59 cell types across 4 lineage classes (∼250 mean counts/cell). **(C) Experimental timecourse.** Lung tissue was sampled at homeostasis (D0) and three post-pneumonectomy timepoints (D3, D7, D14) across n = 3 biological replicates (M1, M2, F1). **(D) Histologic survey across regeneration phases.** Top row: representative whole-lobe H&E sections of the accessory lobe at D0, D3, D7, and D14. Colored boxes mark zoomed locations across four canonical tissue zones: subpleural (red), conducting airway (yellow), parenchyma (brown), and conducting vessel (blue). Below: 100-µm zooms across phases (columns: Homeostasis - D0, Inflammation - D3, Remodeling - D7, Maturation - D14) and zones (rows: Parenchyma, Conducting Airway, Subpleural, Conducting Vessel). Phase-defining histology is shown: D3 mononuclear infiltration of adventitia, sub-pleural space, and alveoli with mesothelial cuboidalization; D7 matrix-dense, microvasculature-sparse foci within parenchyma, sub-pleural space, and adventitia; D14 secondary-septation rings consistent with late-stage alveologenesis and pleural thinning. Scale bars: 2 mm (whole-lobe), 100 µm (zooms). **(E) Cell-type compositional dynamics across regeneration in both modalities.** Stacked bar charts show fractional composition at all timepoints in scRNAseq (left, “sc-Transcriptomics”) and Xenium (right, “sc-ST”), displayed at the lineage-class level (top row, “Class”) and decomposed by Type.6 annotation within each class (Epithelium, Mesenchyme, Immune, Endothelium rows). Both modalities capture concordant inflammatory expansion by D3, transitional cells emergence at D7, and partial resolution toward homeostatic composition by D14. **(F) Spatial findings overview across timepoints.** Whole-puck Xenium fields at D0, D3, D7, and D14 highlight cell populations central to the regeneration process; all other cell types greyed. Activated Mesothelium (magenta), Interstitial Macrophages (yellow), Transitional Macrophages (cyan), Transitional Fibroblasts (purple), and Transitional Epithelium (green). At D0, transitional populations are minimal. By D3, Interstitial Macrophages and Activated Mesothelium accumulate at the subpleural margin. At D7, the three transitional populations co-localize in discrete deep- parenchymal patches. By D14, transitional populations have largely resolved.

Following QC, clustering, and hierarchical annotation, the resulting spatiotemporal atlas resolved major canonical cell types known from human and mouse biology (Fig 1e and Fig S2-S5). Captured epithelium included canonical proximal conducting (Ciliated, Secretory, Basal) and distal subtypes (ATI, ATII) as well as a distinct population of bronchioalveolar stem cells (BASCs). A prominent population of transitional epithelium (Shen et al., 2022; Shin et al., 2025) was also captured on both scRNAseq and Xenium. Subcluster analysis revealed large portions of this transitional population to display clear gene expression similarity with previously-described PATS/DATP/Transitional/Aberrant Basaloid cells (Fig S6) (Choi et al., 2020; Kobayashi et al., 2020; Strunz et al., 2020). Two small subpopulations displayed genes signatures associated with alveolar epithelial progenitors (AEPs) (Zacharias et al., 2018) and activated or proliferating ATII cells (p-ATIIs) (Aspal and Zemans, 2020) (Fig S6).

Beyond epithelium, the atlas captured endothelial (aCaps, gCaps, arterial and venous endothelium, peribronchial endothelium, lymphatic cells), mesenchymal (adventitial and alveolar fibroblasts, distal myofibroblasts, airway and vascular smooth muscle cells, alveolar and peribronchial pericytes, pleural mesothelium), myeloid populations (interstitial and alveolar macrophages, monocytes, neutrophils, mast cells, basophils), and lymphoid populations (T and B cells, Innate Lymphoid Cells (ILCs), plasma cells), with canonical marker expression portrayed in Fig S2-S5. This spatiotemporal atlas, spanning four timepoints, captured both canonical lung populations and a prominent transitional epithelium, providing the substrate for the architectural analyses that follow. A preview of macroscopically distinct cellular ensembles that emerge in our dataset is displayed in Fig1F.

### 2. A multi-layered framework reveals tissue architectural signatures

Close inspection of the spatial transcriptomics data suggested that specific neighborhood compositions are associated with local tissue remodeling. At D3, the subpleural area was observed to be enriched for a unique activated mesothelial population expressing the neutrophilic granulocyte chemoattractant Cxcl6, closely adjacent to a dense concentration of subpleural interstitial macrophages expressing matrix metalloproteases such as Mmp2 (Fig 2a-b). By D7, regions of terminal alveoli were shown to lose broad ATI and ATII predominance and instead were encompassed mainly by a transitional triad of Krt17+ transitional epithelium, Cthrc1+ transitional fibroblasts, and Spp1+ transitional macrophages (Fig 2c-d). D14 showed a morphology consistent with secondary septation, with contractile myofibroblasts emerging at the edges of acinar bulbs (Fig 2e-f).

**Figure 2.**
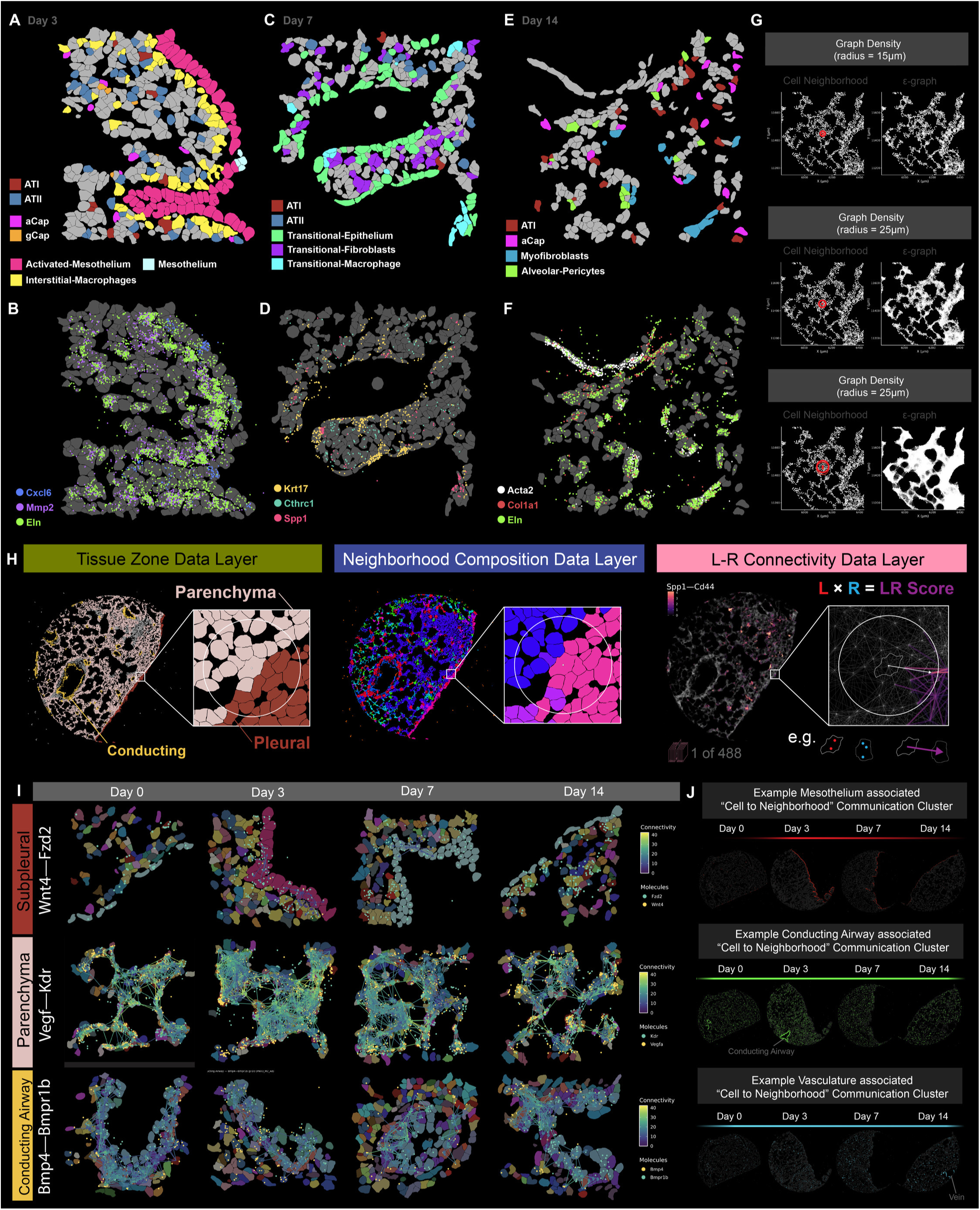
PPLR tissue microenvironments display co-localized signatures of development, regeneration, and disease, revealed across complementary spatial data layers. **(A) Day 3 - Subpleural inflammation microenvironment.** Highlighted cell types include Activated-Mesothelium (magenta) and Mesothelium (pale cyan) at the pleural boundary, Interstitial-Macrophages (yellow) at the subpleural front, and homeostatic parenchymal partners ATI (firebrick), ATII (steel blue), aCap (magenta - purple), and gCap (orange). Other cell types greyed. **(B) Day 3 - Leading-edge gene program.** *Cxcl6* (blue: chemokine recruitment), *Mmp2* (purple: matrix breakdown), and *Eln* (green: developmentally restricted matrix production) co-localize within the subpleural inflammation zone, recapitulating the “matrix breakdown + production” coordination characteristic of developmental and regenerative biology. **(C) Day 7 - Transitional triad in alveolar crest.** Highlighted: Transitional-Epithelium (green), Transitional-Fibroblasts (purple), and Transitional-Macrophages (cyan) co-localizing in a discrete tissue patch flanked by ATI (firebrick) and ATII (steel blue) context. **(D) Day 7 - Transitional triad gene markers.** *Krt17* (yellow: TransEpi), *Cthrc1* (cyan: TransFib), *Spp1* (pink: TransMac). All three transcripts concentrate within the same peninsula, recapitulating the molecular signature of fibroblastic foci described in idiopathic pulmonary fibrosis (IPF). **(E) Day 14 - Secondary septation context.** Highlighted: Myofibroblasts (cyan), Alveolar- Pericytes (lawn green), aCap (magenta), and ATI (firebrick). **(F) Day 14 - Matrix-maturation gene program.** *Acta2* (white: contractile, shared with vascular SMC), *Col1a1* (red: collagen I deposition), *Eln* (green: elastin maturation). The *Acta2* signal aligns with anatomical SMC reference structures (large airway/vessel walls), while *Col1a1* and *Eln* distribute through alveolar septa suggesting active alveologenesis. **(G) Tissue graph construction using three radii.** Each tissue puck is formalized as an undirected radial-neighbor graph; insets show neighborhood definitions at radius = 15 µm (sparse), 25 µm (chosen, recovers ∼10 nearest neighbors per cell on average), and 35 µm (over- merged) for the same reference cell. The 25-µm graph is used for all downstream layers. **(H) Multi-layer spatial-data framework.** Three independently-computed data layers are illustrated on the same tissue: **Tissue Zone** (Parenchyma, Conducting, Pleural; cells colored by territory.coarse), **Neighborhood Composition** (cells colored by neighborhoodPCA.SNN cluster), and **L-R Connectivity** (illustrated for *Spp1*—*Cd44*, with edges drawn between sender and receiver cells weighted by LR score; depicted is one of the 488 ligand-receptor mechanisms scored on every radial-graph edge using NICHES). **(I) Zone-characteristic LR mechanism arcs across timepoints.** Three zone-locked LRMs imaged on matched exemplar pucks across D0, D3, D7, D14. **Subpleural** *Wnt4*—*Fzd2*, **Parenchyma** *Vegfa*—*Kdr*, and **Conducting Airway** *Bmp4*—*Bmpr1b*. Cell polygons colored by Type.6; edge color = LR connectivity score (per-row global viridis scale). Each row traces a distinct temporal arc: Pleural *Wnt4*—*Fzd2* spikes at D3 and resolves by D14; parenchymal *Vegfa*—*Kdr* shows a U-shape with a D7 trough and partial D14 recovery; conducting *Bmp4*— *Bmpr1b* dips at D7 and recovers by D14, establishing that each tissue zone has its own temporal LR signature, all completed by D14. **(J) Example cell-to-neighborhood communication clusters across regeneration.** Local aggregation and clustering of LR-connectivity profiles around each cell reveals signaling patterns specific to cell type and zone communication. Three representative cluster trajectories are shown across D0, D3, D7, D14: a **mesothelium-associated** cluster (top row), a **conducting-airway- associated** cluster (middle row), and a **vasculature-associated** cluster (bottom row). Each cluster waxes and wanes in distinct timepoints, reflecting unique cell-to-neighborhood signaling behaviors tied to the different tissue contexts.

Several proximity-based analyses were designed to allow for statistical analysis of tissue zones, neighborhood composition, and cell-to-cell connectivity. Each tissue sample was first formalized as a graph with cell centroids as nodes and edges connecting nodes only when pairs of cells fell within 25µm of each other (Fig 2g, Fig S7). Three analytic layers were then constructed on this graph. First, well-differentiated landmark cells (such as mesothelial cells at the outer boundary) were used to segregate each cell into specific anatomical tissue zones recognizable as gross anatomical regions such as the pleural border (Fig 2h, left). Second, a ‘neighborhood composition’ assay was created, allowing unbiased clustering of cells based on similarity of local cellular composition (Fig 2h, right). Third, 488 distinct ligand-receptor connectivity layers were mapped on top of the tissue graph using NICHES (Fig 2h, Fig S8; (Raredon *et al*., 2023), stored as a cell-cell signaling ‘hyper-graph’ from which any cell-cell signaling modality captured in our dataset could be visualized spatially. This data structure allowed identification and visualization of the cellular circuits that activated or deactivated over time and space for selected tissue regions (Fig S8). Representative examples of these dynamic connectivity patterns are displayed in Fig 2i. Local aggregation and unbiased clustering of this extensive connectivity data allowed identification of specific clusters of similar “cell-to-neighborhood” signaling associated with cell type transitions (Fig 2j).

This multi-layered framework resolved tissue architecture at three complementary scales: landmark zones identified broad anatomical territories familiar to histologists such as pleural, parenchymal, and conducting airway zones; neighborhood composition captured how cells within these broader zones adopt traditional or unique architectures during response to the pneumonectomy perturbation; and ligand-receptor connectivity characterized the signaling relations between cells rather than treating them as static, disconnected elements that happen to colocalize. These three layers structure the section-by-section analyses of D3, D7, and D14 that follow.

### 3. A boundary-activation architectural signature predominates at D3

The subpleural community emerging at D3 (Fig 3a-b) was characterized by mesothelial cells undergoing a shift in cell state, upregulating ligands spanning neutrophil chemoattraction (Cxcl6), vascular-stress signaling (Angptl4), and FGF mitogenesis (Fgf1) on both our single cell and spatial analyses (Fig 3c). This mesothelial activation coincided in space and time with dense subpleural congregations of canonical interstitial macrophages (Fig 3a-c). Both Yap1 (Fig 3d-e) and downstream mechanosensitive channels including both Piezo1/2 and Ccn2 (Ctgf) were observed to be prominently enriched in both the pleura and parenchyma of the lung at D3 (Fig S9). By D3, parenchymal cell-type neighborhoods were significantly enriched for both interstitial and alveolar macrophages (Fig S10), which co-localized in particular with ATI and ATII cells, alveolar fibroblasts, and alveolar endothelium.

**Figure 3.**
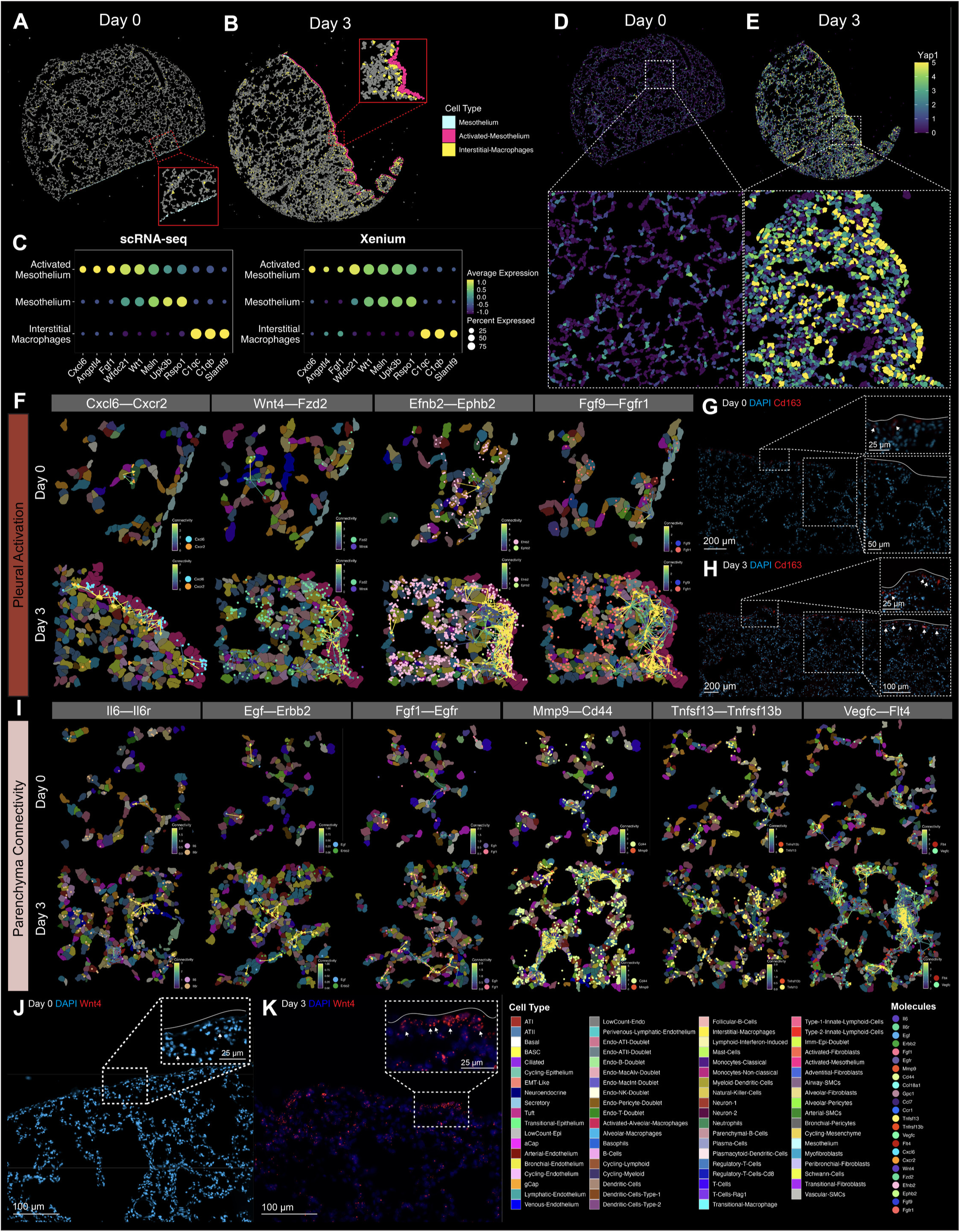
Pleural boundary activation initiates regeneration at Day 3 and signaling changes propagate into the parenchyma. **(A) Day 0 - Subpleural cellular community.** Whole-puck highlighting Mesothelium (pale cyan), Activated-Mesothelium (magenta), and Interstitial-Macrophages (yellow); other cell types greyed. Mesothelium at D0 is restricted to a thin pleural layer with sparse interstitial macrophages distributed through the parenchyma. **(B) Day 3 - Subpleural cellular community.** Same color scheme as (A). At D3, Activated- Mesothelium is localized thorough the entire pleural edge and accumulated subpleural Interstitial- Macrophages line the boundary. **(C) Marker expression of pleural-zone cell types in scRNAseq and Xenium.** Dotplot of canonical and induced markers across Activated-Mesothelium, Mesothelium, and Interstitial- Macrophages, in scRNAseq (left) and Xenium (right). Activated-Mesothelium is defined by induced expression of *Cxcl6*, *Angptl4*, *Fgf1*, *Wfdc21* alongside conserved mesothelial markers (*Wt1*, *Msln*, *Upk3b*, *Rspo1*); Interstitial-Macrophages defined by *C1qc*, *C1qb*, *Slamf9*. Both modalities show concordant marker patterns, supporting cell-type assignments. Dot size = fraction of cells expressing; color = average expression (z-scored). **(D) Day 0 - *Yap1* spatial map.** *Yap1* transcript counts per cell (viridis: 0–5+). During homeostasis, *Yap1* is low and uniform across the tissue. **(E) Day 3 - *Yap1* spatial map.** At D3, *Yap1* expression is broadly induced, especially concentrating at the pleural margin and along emerging activated parenchymal edges. *Yap1* induction parallels mechanical-stress-driven nuclear translocation in development and injury contexts; companion *Piezo1*/*Piezo2* maps appear in Fig S6. **(F) Pleural-zone single-edge L-R connectivity changes from Day 0 to Day 3.** Spatial connectivity plots show four pleural-axis LR mechanisms in matched crops at D0 (top row) vs D3 (bottom row): *Cxcl6*-*Cxcr2* (chemokine-mediated myeloid recruitment), *Wnt4*-*Fzd2* (Wnt- dependent epithelial/mesothelial state transition), *Efnb2*-*Ephb2* (Eph-Ephrin boundary and contact-guidance signaling), *Fgf9*-*Fgfr1* (FGF-driven branching morphogenesis and repair). Cells colored by Type.6; ligand and receptor transcripts shown as labeled dots; edge color = LR connectivity score (viridis). Across all four mechanisms, connectivity density and edge weight increase substantially from D0 to D3. **(G) Day 0 — Anti-Cd163 immunohistochemistry.** Paraffin-embedded D0 lung sections stained for DAPI (blue) and Cd163 (red, an Interstitial-Macrophage marker). At D0, sparse Cd163+ cells are distributed through parenchyma; the pleural edge shows minimal localization. **(H) Day 3 — Anti-Cd163 immunohistochemistry.** Same staining and scale as (G). At day 3, Cd163+ cells accumulate concentrically along the pleural margin, independently validating the recruitment pattern in (B). **(I) Parenchymal single-edge L-R connectivity changes from Day 0 to Day 3.** Spatial connectivity plots of six parenchymal-axis LRMs at D0 (top row) vs D3 (bottom row): *Il6*-*Il6r* (inflammatory), *Egf*-*Erbb2* (epithelial growth), *Fgf1*-*Egfr* (cross-family epithelial growth), *Mmp9*- *Cd44* (matrix remodeling), *Tnfsf13*-*Tnfrsf13b* (immune-stromal crosstalk), *Vegfc*-*Flt4* (lymphangiogenic and vascular regeneration). All six mechanisms show increased intra- parenchymal signaling density at D3, consistent with pleural activation propagating inward into a broad parenchymal response. **(J) Day 0 - *Wnt4* RNAscope FISH.** Detection of *Wnt4* mRNA (red) with DAPI nuclear counterstain (blue). At D0, *Wnt4* transcripts are sparsely detected at the pleural edge. **(K) Day 3 - *Wnt4* RNAscope FISH.** Same staining and scale as (J). At D3, *Wnt4* is markedly elevated specifically along the subpleural boundary, independently validating the LR-connectivity finding for *Wnt4*-*Fzd2* in (F) and the Fig 2(I) pleural row arc.

The connectivity layer resolved which signals were actively exchanged between co-localized cells. We isolated the cell-cell signaling interactome by cell type and found the mesothelial interactome to be the most highly perturbed of all cells profiled within the lung at D3 (Fig S11), with this activation associated with diverse upregulation of inflammatory and developmental connectivity. This included enrichment of neutrophil chemoattraction (Cxcl6-Cxcr2), Wnt-mediated developmental patterning (Wnt4-Fzd2), Eph/ephrin boundary patterning (Efnb2-Ephb2), and FGF mesenchymal growth signaling (Fgf9-Fgfr1) at the pleural border at D3 (Fig 3f).

The subpleural enrichment of transcriptionally defined interstitial macrophages at D3 was independently validated by *in situ* CD163 immunostaining, which demonstrated a dense accumulation of CD163⁺ macrophages beneath the pleura, found also prior (Fabriek et al., 2005; Yamashita et al., 2018). (Fig 3g-h). Beyond the pleural niche, parenchymal connectivity was also highly perturbed at D3, with concurrent elevation of inflammatory (Il6-Il6r), proliferative (Egf-Erbb2, Fgf1-Egfr), matrix-remodeling (Mmp9-Cd44), immunoregulatory (Tnfsf13-Tnfrsf13b), and lymphangiogenic (Vegfc-Flt4) signaling (Fig 3i). Wnt4, a developmental Wnt ligand normally absent during homeostasis (Fig 3j), was confirmed by RNAscope to be broadly upregulated in both the parenchyma and the pleura by Day 3 (Fig 3k). The co-occurrence of mechanosensitive expression, inflammatory infiltration, and developmental cue activation at D3 is consistent with a boundary-initiated, mechano-inflammatory response coordinating the early post-pneumonectomy lung.

### 4. A fibrosis-mimetic transitional architectural signature is enriched at D7

By D7 post-pneumonectomy, regions of the parenchyma showed signs of remodeling that bore striking histologic resemblance to features seen in pulmonary fibrosis (Fig 4a-b). In our spatial transcriptomics data, these regions were not observed at D0, and were only observed minimally at D3 in scattered nucleations. Fibrogenic regions were heterogeneously distributed across replicate animals at D7, and were essentially absent by D14. Whole-accessory-lobe histologic survey of D7 lungs revealed these remodeling regions occurring as multiple discrete foci across the parenchyma; therefore, our Xenium cores captured representative examples of a recurrent tissue state rather than isolated aberrations (Fig S12). The gross morphology of these regions also varied across core samples: some displayed retained parenchymal branching features within the remodeling zone (Fig 4a), while others showed dense cellular consolidation with near-complete loss of alveolar architecture (Fig 4b), suggesting heterogeneity in either the morphological manifestation of this phenomenon or the capture of different stages along its course.

**Figure 4.**
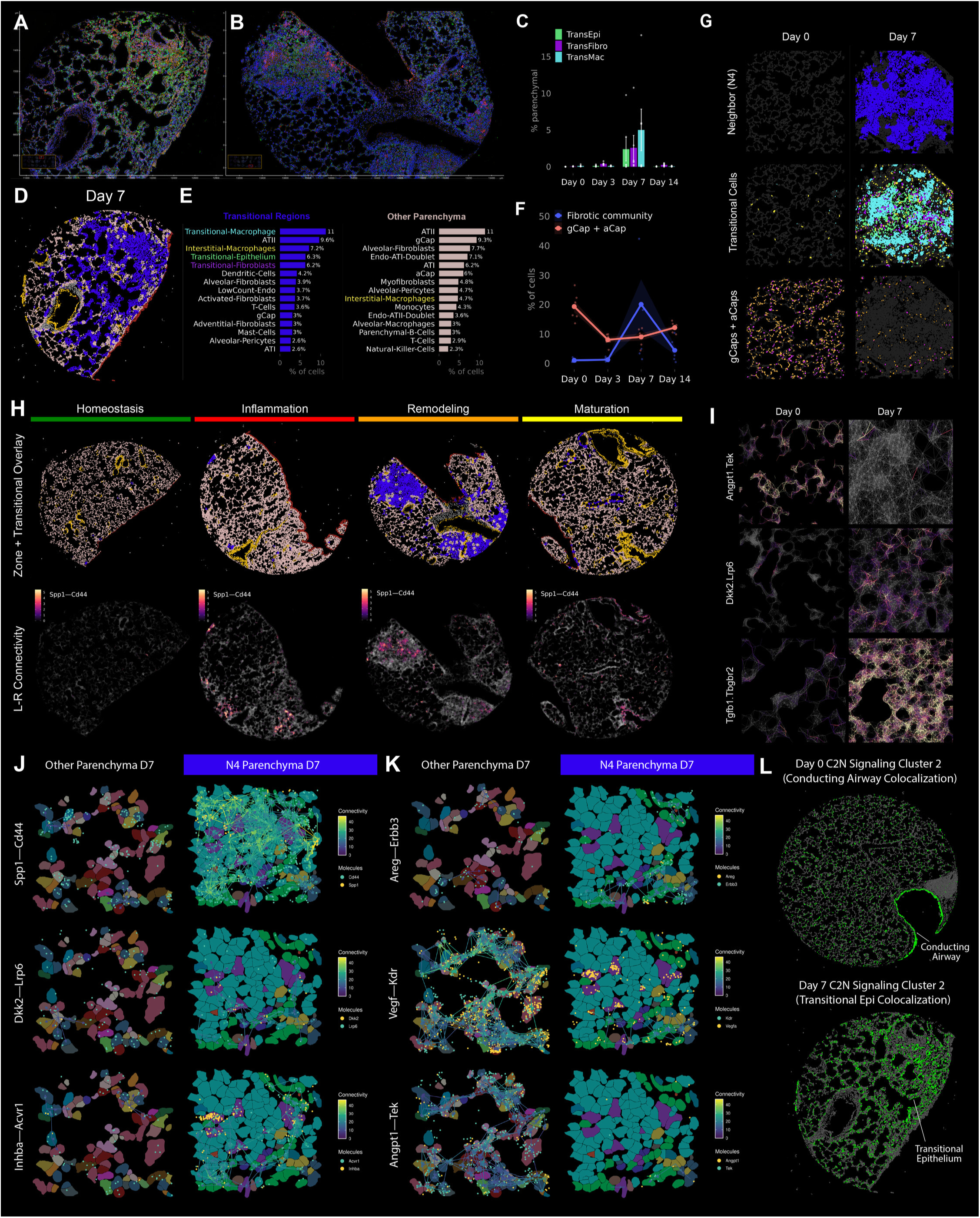
A discrete transitional remodeling community emerges in the deep parenchyma at Day 7, defined by enriched fibrotic-axis and depleted vascular-axis ligand-receptor signaling. **(A) D7 transitional region – full Type.6 spatial overlay.** Cells colored by Type.6 annotation to show the cellular composition of the highlighted transitional patches: Transitional- Epithelium (green), Transitional-Fibroblasts (purple), and Transitional-Macrophages (cyan) co-aggregate within the patches, with Interstitial-Macrophages (yellow) accumulated at patch boundaries. **(B) D7 fibrotic-mimetic region.** Same coloring as (A) by Type.6 annotation. Highlights large dense cellular consolidations of Transitional-Epithelium, Transitional-Fibroblasts, and Transitional-Macrophages, and loss of functional architecture. **(C) Temporal emergence of the transitional triad across regeneration.** Bar chart of Transitional-Epithelium (green), Transitional-Fibroblasts (purple), and Transitional- Macrophages (cyan) as a percent of parenchymal cells per timepoint. All three populations emerge from near-zero at D0/D3, peak at D7 (means 2–5% per type with one puck outlier reaching ∼16%), and resolve to baseline by D14. Bars: mean ± SD across pucks; dots: per-puck values. **(D) D7 fibrotic-density spatial overview.** Whole-puck spatial plot showing fibrotic-density labeling per cell on a representative D7 sample: cells assigned to the transitional / fibrotic- mimetic community are highlighted in blue, conducting structures in yellow, all other cells grey. Discrete coherent patches of transitional community are visible in deep parenchyma, sparing the subpleural margin. **(E) Cell composition of D7 transitional regions vs other parenchyma.** Top 15 cell types by parenchymal-cell fraction in matched D7 regions. **N4 / Transitional regions** (left, blue bars) are enriched for the canonical triad - Transitional-Macrophage, Transitional- Epithelium, Transitional-Fibroblasts - alongside ATII and Interstitial-Macrophages. **Other parenchyma** (right, cream bars) shows the homeostatic distribution dominated by ATII, gCap, Alveolar-Fibroblasts, Endo-ATI-Doublet, and ATI. Triad populations are essentially absent from non-transitional parenchyma; gCap is essentially absent from transitional regions. **(F) Reciprocal temporal dynamics of fibrotic community vs alveolar capillaries.** Line plot across timepoints, mean ± SD across pucks. **gCap + aCap fraction** (coral) starts high at D0, drops sharply at D3, reaches a nadir at D7, and recovers partially by D14. **Fibrotic- community fraction** (blue) is near-zero at D0/D3, peaks at D7, and resolves by D14. The two trajectories are temporally inversely correlated, consistent with the spatial co- emergence shown in (G). **(G) Spatial co-emergence of N4 cluster, transitional cells, and capillary loss at D7.** Three rows compare D0 (left) vs D7 (right) on the same exemplar puck. **Top - Neighbor (N4):** the N4 neighborhood cluster (blue) is essentially absent at D0 and dominates parenchymal regions at D7. **Middle - Transitional cells:** Transitional-Epithelium (green), Transitional-Fibroblasts (purple), and Transitional-Macrophages (cyan) co-aggregate within the same D7 regions; sparse at D0. **Bottom - gCaps + aCaps:** capillary populations (orange and magenta) are notably depleted from regions where transitional cells aggregate at D7 (D0 baseline at left for reference). **(H) Tissue zone and transitional overlay (top) and *Spp1*-*Cd44* connectivity (bottom) across regeneration timepoints.** Both rows show the same exemplar puck across Homeostasis (D0), Inflammation (D3), Remodeling (D7), and Maturation (D14). **Top:** tissue zones (Parenchyma in cream, Pleura in red, and Conducting Airways in yellow) overlaid with the Transitional / N4 cluster (blue) -visible only at Remodeling and absent at Maturation. **Bottom:** *Spp1*-*Cd44* LR connectivity (color = edge weight) on the same pucks, peaking at Inflammation/Remodeling and declining at Maturation, illustrating how a single LRM’s temporal arc tracks the emergence-and-resolution of the transitional cluster. **(I) Three contrasting LR mechanisms at Day 0 vs Day 7 in matched parenchymal crops.** *Angpt1*-*Tek* (vessel maturation): dense connectivity at D0 collapses to sparse residual edges at D7. *Dkk2*-*Lrp6* (WNT antagonism) and *Tgfb1*-*Tgfbr2* (TGF-β axis): opposite pattern - sparse at D0, dense at D7. Together, at peak remodeling, vascular maintenance signaling falls while fibrotic-axis signaling rises within the same parenchymal regions. **(J) Direct N4 vs non-N4 D7 LR comparison: fibrotic-axis enrichments and WNT antagonism.** Three locked LR mechanisms imaged in matched D7 transitional (“N4 Parenchyma D7”) and non-transitional (“Other Parenchyma D7”) regions of equal area (∼13,500 µm²): *Spp1*-*Cd44* (fibrotic-macrophage / fibroblast hallmark recapitulating the IPF SPP1+/CD44+ axis), *Dkk2*-*Lrp6* (WNT antagonism, blocking AT2→AT1 differentiation), and *Inhba*-*Acvr1* (Activin-A / ADI signature). All three are enriched in N4 Parenchyma at D7. Cells colored by Type.6; ligand and receptor transcripts shown as labeled dots; edge color = LR connectivity score (global viridis scale shared with K). **(K) Direct N4 vs non-N4 D7 LR comparison: repair-attempt cue and depleted vascular- axis mechanisms.** Three additional LR mechanisms imaged in the same matched D7 regions as (J): *Areg*-*Erbb3* (amphiregulin-driven repair-attempt cue, enriched in N4), *Vegfa*-*Kdr* (vascular induction, depleted in N4), and *Angpt1*-*Tek* (vessel maturation, depleted in N4). Notably, *Vegfd*-mediated connectivity is also depleted in N4 (Fig S4). The N4 LR signature is thus dominated as much by what is *lost* (vascular induction, vessel maturation, and barrier protection) as by what is *gained* (fibrotic-mimetic, WNT-antagonist, ADI, and repair-attempt cues), consistent with a conditionally-arrested state induced by local endothelial deficit. **(L) Cell-to-neighborhood signaling cluster examples across phases.** Two illustrative cell- to-neighborhood communication clusters: **Day 0 - C2N Signaling Cluster 2** showing Conducting Airway Colocalization at homeostasis, and **Day 7 - C2N Signaling Cluster 2** showing Transitional Epithelium Colocalization at peak remodeling.

Cellular composition of these tissue regions at D7 was dominated by a co-aggregating triad: Krt17+ transitional epithelium, Cthrc1+/Dkk2+ transitional fibroblasts, and Spp1+ transitional macrophages, accompanied by a moderate enrichment of interstitial macrophages and a marked depletion of well-differentiated alveolar capillary populations (gCaps+aCaps) (Fig 4C-E). All three transitional populations were rare at D0 and D3, enriched at D7, and near baseline by D14: a pattern consistent with transient emergence and resolution (Fig 4C). Unbiased neighborhood clustering of local cellular composition identified a distinct cluster, labeled here as Neighborhood 4 (N4), that was enriched for this transitional triad and depleted for capillary populations (Fig 4D-E). Spatial co-emergence at D7 was evident: the N4 region was observed to dominate parenchyma in samples at D7 and was essentially absent at D0 (Fig 4F-G). Investigation of population-level temporal dynamics confirmed the reciprocal relationship between the fibrotic-community fraction and the capillary (gCap+aCap) fraction. Capillary cell populations reached their nadir while transitional fractions reached their peak simultaneously at D7, and both partially returned toward homeostatic levels by D14 (Fig 4F-G).

The N4 region displayed a high-dimensional connectivity signature, with elevated connectivity mediated by TGFβ-superfamily signaling (Tgfb1, Tgfb2, Inhba), EGF-family epithelial cues (Areg, Ereg), developmental Wnt patterning (Wnt7b), matrix scaffolding (Spp1, Lama5/b3/c2), and chemokines and cytokines (Cxcl16, Il24) (Fig 4H, Fig S13). N4 parenchyma at D7 showed a loss of vessel-maturation connectivity (Angpt1-Tek) alongside marked gains in Wnt antagonism (Dkk2-Lrp6) and pro-fibrotic TGFβ signaling (Tgfb1-Tgfbr2) as compared to homeostatic control regions (Fig 4I). N4 parenchyma showed significant loss of endothelial-homeostasis connectivity, including endothelial maintenance (Apln-Aplnr), VEGF angiogenesis (Vegfa-Kdr; depleted alongside Vegfa-Flt1 and Vegfa-Nrp1), and vessel maturation (Angpt1-Tek) (Fig S13).

This connectivity signature was found to be a property of the transitional tissue state itself (N4), not of D7 parenchyma in aggregate. To establish this, matched N4-rich and non-transitional regions of equal area were directly compared within the same D7 samples (Fig 4J-K). This analysis revealed matrix-remodeling adhesion (Spp1-Cd44), Wnt antagonism (Dkk2-Lrp6), activin signaling (Inhba-Acvr1), and EGF-family proliferation (Areg-Erbb3) enriched within transitional regions, while VEGF angiogenesis (Vegfa-Kdr) and vessel maturation (Angpt1-Tek) were depleted.

Independent unsupervised clustering of cell-to-neighborhood signaling profiles further revealed a communication cluster that co-localized with the conducting airway at all timepoints but also co-localized with transitional epithelium throughout the parenchyma at D7 (Fig 4L). This finding is consistent with the described phenomenon of “proximalization”, the adoption of proximal airway-like signaling character by distal transitional epithelium (Jaeger et al., 2022; Tata and Rajagopal, 2017; Yang et al., 2017). The transitional epithelium’s heterogeneous distribution at D7 and near-absence at D14 is consistent with transient enrichment during regeneration rather than stable pathology.

### 5. An alveologenic architectural signature characterizes D14

By D14 post-pneumonectomy, the distal acini at the ends of the airway tree showed histologic signs of alveologenic maturation (Fig 1d, Fig S1), with secondary-crest myofibroblasts appearing within alveolar ring cross-sections, co-localized with well-differentiated distal epithelial, endothelial, and mural populations (Fig 2e-f). Over our time course, the thin septa and intact mesothelial boundary seen at homeostasis gave way to border hypercellularity at D3, thickened primary septa at D7, and by D14 we observe secondary septa bearing secondary-crest myofibroblasts at their tips (Fig 5A).

**Figure 5.**
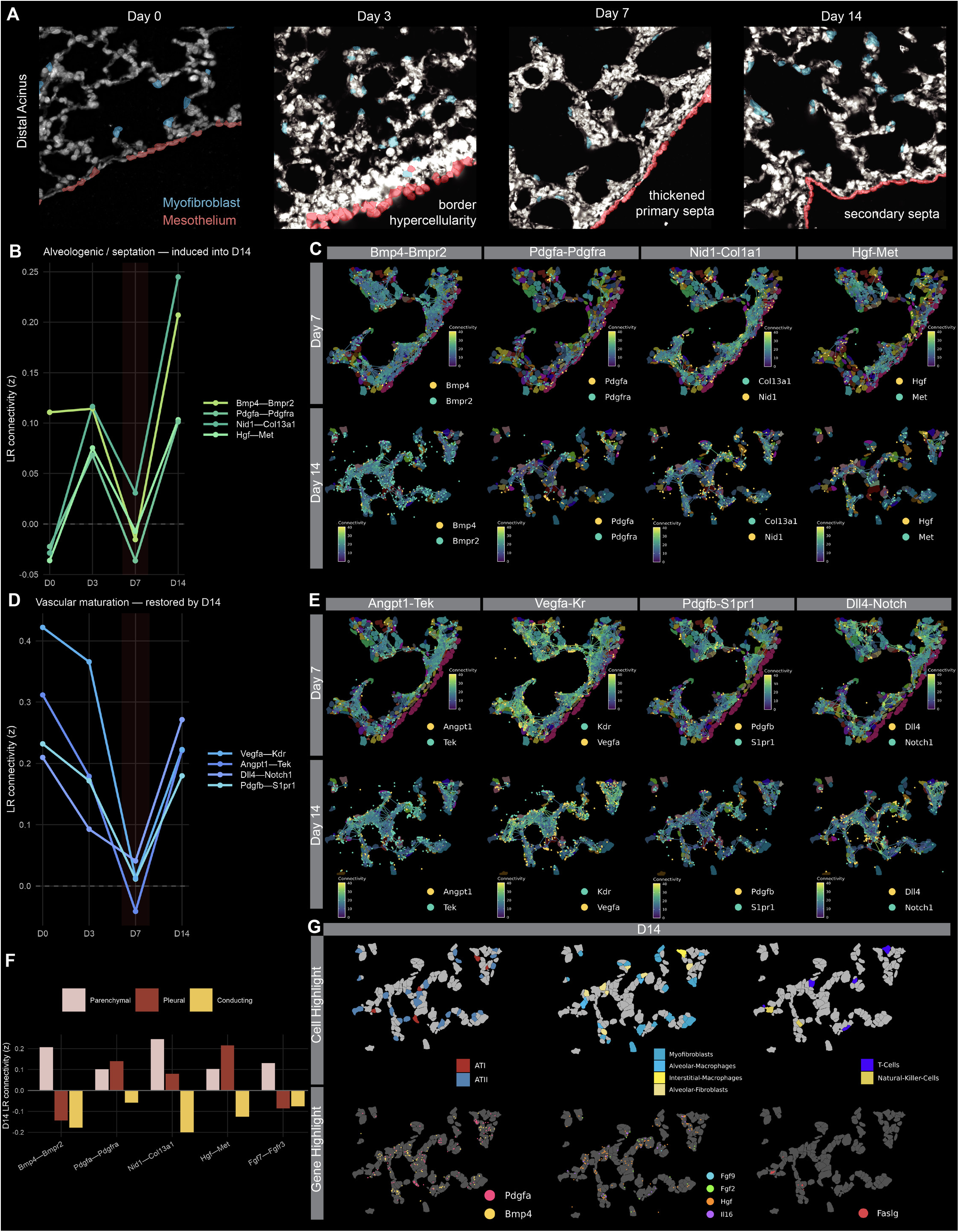
Day 14 architecture is defined by an induced alveologenic signaling program and a restored vascular program. (A) Representative spatial cell maps of the distal acinus across the regeneration time course (Day 0, 3, 7, 14), with myofibroblasts (blue) and mesothelium (red) highlighted; all other cells grey. Annotated histologic features: border hypercellularity at D3, thickened primary septa at D7, and nascent secondary septa at D14. (B) Temporal trajectories of aggregate parenchymal ligand–receptor (LR) connectivity (z-score) across D0–D14 for four alveologenic/septation mechanisms (Bmp4–Bmpr2, Pdgfa–Pdgfra, Nid1–Col13a1, Hgf–Met), each reaching maximal connectivity at D14. (C) Spatial LR connectivity plots for the four mechanisms in (B) at Day 7 and Day 14. Pucks: D7, [PNX7_M1_Sp]; D14, [PNX14_M2_Ad]. (D) Temporal trajectories of aggregate parenchymal LR connectivity (z-score) for four vascular-maturation mechanisms (Vegfa–Kdr, Angpt1–Tek, Dll4–Notch1, Pdgfb–S1pr1), elevated at baseline, lowest at D7, and recovering toward baseline by D14. (E) Spatial LR connectivity plots for the four mechanisms in (D) at Day 7 and Day 14, displayed as in (C). (F) D14 LR connectivity (z-score) across the parenchymal, pleural, and conducting landmark zones for five alveologenic mechanisms (Bmp4–Bmpr2, Pdgfa–Pdgfra, Nid1– Col13a1, Hgf–Met, Fgf7–Fgfr3), enriched predominantly within the parenchymal acinus. (G) In situ localization of D14 ligands to their cellular sources in a representative D14 acinus ([PNX14_M2_Ad]). *Top (cell highlight):* cell types of interest in MSBR colors, all others grey — (left) ATI, ATII; (middle) myofibroblasts, alveolar macrophages, interstitial macrophages, alveolar fibroblasts; (right) T cells, natural-killer cells. *Bottom (gene highlight):* corresponding ligand transcripts as dots on grey cells — (left) Pdgfa, Bmp4; (middle) Fgf9, Fgf2, Hgf, Il16; (right) Faslg.

Tissue maturation at D14 displayed a unique connectivity signature not found within other tissue phases. Most signaling mechanisms perturbed earlier in the time course returned within their homeostatic baseline at D14 (Ccl2, Cthrc1, Cxcl6, Dkk2, Fn1, Il17b, Lama5, Lamc1/2, Ntf4, Ntn3, Slit2, Spp1, Tgfb1, Tnc, Wnt7b), and most remaining mechanisms reverted toward D3-like inflammatory levels, consistent with ongoing activation and incomplete return to homeostasis. Two signaling programs stood out as elevated at D14 relative to every other phase, including baseline.

The first was a set of mechanisms that can be considered to comprise an alveologenic program at D14. A set of mesenchymal-mitogen and morphogen mechanisms (Bmp4-Bmpr2, Pdgfa-Pdgfra, Nid1-Col13a1, and Hgf-Met) reached their maximal connectivity at D14, exceeding baseline and all intervening timepoints (Fig 5B), and LR visualization of distal acini showed these edges emerging across the parenchyma between remodeling D7 and D14 parenchyma (Fig 5C). Several of these mechanisms have been associated with the secondary-crest myofibroblast and its epithelial neighbors. Pdgfa-Pdgfra signaling has been linked to myofibroblast positioning at the septal tip, Bmp4 and Hgf to alveolar morphogenic and epithelial cues, and Nid1-Col13a1 to basement-membrane adhesion (a broader set of D14-elevated signals also included mesenchymal-mitogen and morphogen signaling through Fgf2/Fgf9, Wnt potentiation through Rspo1/2, death-receptor ligand signaling through Faslg/Tnfsf10, and immune cytokine signaling Il15) (Fig 5C). Most of these signals were enriched specifically within the parenchymal acinus rather than the pleural or conducting-airway zones, though notably Pdgfa-Pdgfra and Hgf-Met also showed pleural enrichment, while Bmp4-Bmpr2 and Nid1-Col13a1 were the most parenchyma-restricted (Fig 5F).

The second set of signaling programs related to maturing vascular signaling. The angiogenic and vessel-maturation mechanisms Vegfa-Kdr, Angpt1-Tek, Dll4-Notch1, and Pdgfb-S1pr1 were elevated at baseline, fell to their lowest connectivity at D7 (coincident with the microvascular rarefaction of the N4 fibrosis-mimetic state) and recovered toward baseline by D14 (Fig 5D-E). Localizing D14 ligands to their cellular sources, we observed each transcript concentrated over a distinct cellular compartment (Fig 5G). Within the alveolar epithelium, Bmp4 transcripts were concentrated over ATII cells and Pdgfa over ATI cells. The mesenchymal and myeloid cues separated by source, where Fgf9 and Fgf2 localized to myofibroblasts, Hgf localized to alveolar and interstitial macrophages, Il16 to interstitial macrophages and alveolar fibroblasts, and the death-receptor ligand Faslg localized to T cells. The alveologenic architecture at D14 is thus defined by active morphogenic signaling, not just by return to homeostasis. Where D7 carries a fibrosis-mimetic signature consistent with resolution, D14 carries a connectivity signature consistent with active de novo growth and maturation distinct from any other timepoint in the dataset.

## Discussion

In our four-timepoint atlas, we read our findings as the coordinated interplay of agents (cells) within a material (tissue), each balancing its own homeostasis against the collective’s to cause the material to traverse an array of stable points. Here, we deliberately pushed the lung far from its homeostatic baseline, removing an entire left lobe and forcing the remainder to grow to a new size, and asked, in space and time, how cellular agents within the material collectively resolve the perturbation (Fig6A-6B). Pneumonectomy is a broad perturbation in the tissue-systems sense, and read at single-cell and spatial resolution, the response was not uniform over time. Three distinct architectures emerged in this heterogeneously adapting tissue, each dominating a successive timepoint: boundary activation at D3, transitional fibrosis-mimesis at D7, and alveologenesis at D14 (Fig6C)

We interpret these three architectures as successive positions along a continuum of adaptation and resistance (Fig6D-E). The transient and stable configurations the tissue adopts as it works back toward homeostasis are set by each one’s position in perturbational space, by the magnitude of the local perturbation, and by whether endogenous effectors can still adapt to it. Despite their distinct cellular compositions and spatial organization, each architecture draws upon the same broad classes of response the tissue can mount, a mechanical response, an inflammatory response, and a redeployment of developmental signaling, with a different combination dominant at each timepoint. Mechanotransduction, inflammation, and developmental coupling define D3; proximalization, fibrosis-mimetic remodeling, and microvascular rarefaction define D7; and a distinct alveologenic morphogenic signature defines D14. Together, these timepoints trace the canonical arc of regenerative programs profiled across multiple organ systems, from an inflammatory exit from homeostasis, through reactivation of developmental patterning, to tissue maturation. This coupling of mechanotransduction, inflammation, and developmental reprogramming recurs across digit, liver, kidney, and skin (Dolan et al., 2022; Grzelak et al., 2023; Sun et al., 2022; Ye et al., 2024), with HIF signaling proposed as a general axis given the tight interweaving of parenchyma with vasculature (DeFrates et al., 2021).

Why do we see discrete regional signatures rather than a uniform whole-organ response? One explanation is that local differences in perturbation and response magnitudes give rise to distinct stable tissue states. A tissue spanning a range of local conditions can settle into a small number of stable configurations instead of varying smoothly (Jaeger and Monk, 2014). The spatial nucleation pattern of the N4 remodeling region across our dataset is consistent with this behavior.

The lung’s anatomy also constrains the cellular ensembles because the pliability of the organ, and therefore its strain, varies systematically by position. As a branching, arborized organ enclosed by the visceral pleura, the lung is unlikely to distribute post-pneumonectomy strain uniformly across its airways and lobes. Mechanical heterogeneity extends beyond the branching geometry itself. The visceral pleura buckles under its own growth dynamics, while the parenchyma holds alveolar epithelium and microvasculature in near-direct apposition and exhibits a proximal-to-distal rigidity gradient (Hsia, 2017).

Position-specific mechanotransduction thus offers one unifying explanation for the regionalized signaling patterns that we observed in the regenerating lung, including pleural Wnt4-Fzd2, distal-parenchymal Vegfa-Kdr, and conducting-airway Bmp4-Bmpr1b (Figure 2I): positional geometry is associated with variable biomechanics that therefore predicts non-uniform activation. During development, SOX2 and SOX9 partition proximal from distal epithelial fate while the FGF, SHH, Wnt, VEGF, and PDGF families coordinate epithelium and mesenchyme as the airway tree branches (Warburton, 2021), and the reappearance of many of these signals during PPLR is plausibly the reactivation of some of these same morphogenic programs.

The lung’s central mechanical challenge after pneumonectomy is not simply to enlarge existing alveoli, but to enlarge the organ itself through coordinated growth of the airway and vascular trees. While alveolar enlargement and increased packing density provide an initial compensatory reserve, they cannot indefinitely accommodate the addition of new alveolar units. Ultimately, the mesothelial boundary must expand, allowing the lung to physically grow. The D3 architecture shows how position-specific mechanotransduction solves that problem at the boundary. Inflammation engages the entire tissue, but it is apparent most distinctly at the pleural edge, where the mesothelium undergoes a coordinated activation, upregulating neutrophil chemoattraction (Cxcl6), vascular-stress signaling (Angptl4), and FGF mitogenesis (Fgf1) directly above a dense, single-cell-thick row of subpleural interstitial macrophages (Figure 3A-B). The visceral pleura behaves as a boundary regulator of post-pneumonectomy regeneration, paralleling boundary-regulated programs in which a thin epithelial edge governs the patterning and proliferation of underlying mesenchyme, including the apical ectodermal ridge in limb development (Niswander et al., 1993) and the imaginal disc edge in Drosophila (Diaz-Benjumea and Cohen, 1993). Captured in a tissue that subsequently regenerates, this boundary-layered organization appears to be a feature of productive regeneration rather than injury or development alone, and it supplies the mechanical license for growth. Local matrix remodeling at the mesothelial edge by recruited Mmp2+/Eln+ subpleural macrophages could plausibly let the enclosing sac yield both the physical space and the substrate required for newly formed alveolar units.

The shift away from homeostatic identity extends beyond single cells. At D7, transitional identity appears at the level of whole multicellular ensembles. Heterogeneously distributed parenchymal regions at D7 adopt a histology and cellular composition resembling a fibrotic niche, dominated by a co-aggregating triad of Krt17+ transitional epithelium, Cthrc1+ transitional fibroblasts, and Spp1+ transitional macrophages, with marked local depletion of well-differentiated alveolar capillaries (Figure 4C-E). An alveolar unit can transiently retain its architecture despite being populated almost entirely by transitional cell states (Figure 2C, Fig6C). Cellular identity is therefore displaced from homeostasis before tissue architecture is lost. This reframes the question from identifying altered cell states to whether the tissue unit together can re-enter a homeostatic state, which has implications for human disease.

Our PPLR atlas offers the first spatially resolved model of lung regeneration in which both arms of the regeneration-fibrosis dichotomy can be observed and compared in vivo. The transitional epithelial cells at D7 resemble the aberrant basaloid populations of IPF (Adams et al., 2020) and the repair-associated transitional cells described across models of lung injury (Choi *et al*., 2020; Konkimalla *et al*., 2023; Strunz *et al*., 2020), but our spatial data extend the picture. Where prior work identified a co-localizing transitional epithelial-fibroblast pair (Konkimalla *et al*., 2023), our data extend this model with a co-aggregating triad that adds a Spp1+ transitional macrophage state within capillary-depleted regions. The convergence of “proximalized” epithelium, matrix-producing stroma, and capillary depletion has a developmental precedent, as distal airway progenitors transiently occupy matrix-rich, microvasculature-sparse niches before resolving (He et al., 2022; Warburton, 2021). Additionally, the gene programs and signals we see activated in D7 mirror developmental reactivation that has been previously described as implicated in idiopathic pulmonary fibrosis (Ke et al., 2025b; Selman et al., 2008).

We hypothesize, from the pattern across our timepoints, that the D7 state sits at a bifurcation between two stable points: either the remodeling tissue returns toward homeostasis, or it progresses into chronic fibrosis. Comparable bifurcations have been described in macrophage-fibroblast repair circuits, where the same circuit resolves toward healing or tips into fibrosis depending on its cell ratio (Adler et al., 2020). Read through the framework of the ‘minimal tissue unit’, in which a small group of general populations divide labor to maintain local homeostasis (Adler *et al*., 2023), the alveolar parenchyma has four components: epithelium, fibroblast, macrophage, and microvasculature. We find here that during regeneration, three of the four adopt transitional states while the fourth, well-differentiated microvasculature, is locally lost, which positions endothelial loss as a key disrupting parameter. This is a loss of differentiated, gas-exchanging capillary identity, not a loss of endothelial cells themselves, consistent with cross-organ evidence that fibrogenesis involves microvascular rarefaction (Kida et al., 2014; Manetti et al., 2010).

We represent this in Figure 6E-F as a possible excursion off the homeostatic cycle, in which the unit leaves the inflammation-to-maturation segment and stretches its adaptive range, the set of configurations from which return to homeostasis remains possible. The transitional regions were themselves heterogeneous in form, from foci that retained parenchymal branching to others showing dense consolidation and near-complete loss of alveolar architecture (Figure 4a-b). This can be interpreted as tissue units caught at different depths along the same detour, some closer to folding back and others closer to escape. Two fates likely follow from this precipice. The unit can fold back toward resolution, or, under sustained or repeated demand, escapes the adaptive range into the self-reinforcing fibrotic state seen in chronic fibrotic diseases such as IPF and COPD. Which fate it takes depends on whether endogenous repair preserves resolution capacity before additional demand overwhelms it, and in our dataset the D14 trajectory is consistent with the former, as the transitional triad is near-absent. The arrest is therefore conditional, in that the unit is held off the homeostatic path and whether it returns depends on the shared repair-and-fibrosis machinery not being overwhelmed (Henderson et al., 2020). Our four-component depiction is a deliberate simplification, future work will be needed to incorporate additional features such as component ratios, intercellular distances, and the proximity ranges required to maintain tissue stability.

**Figure 6.**
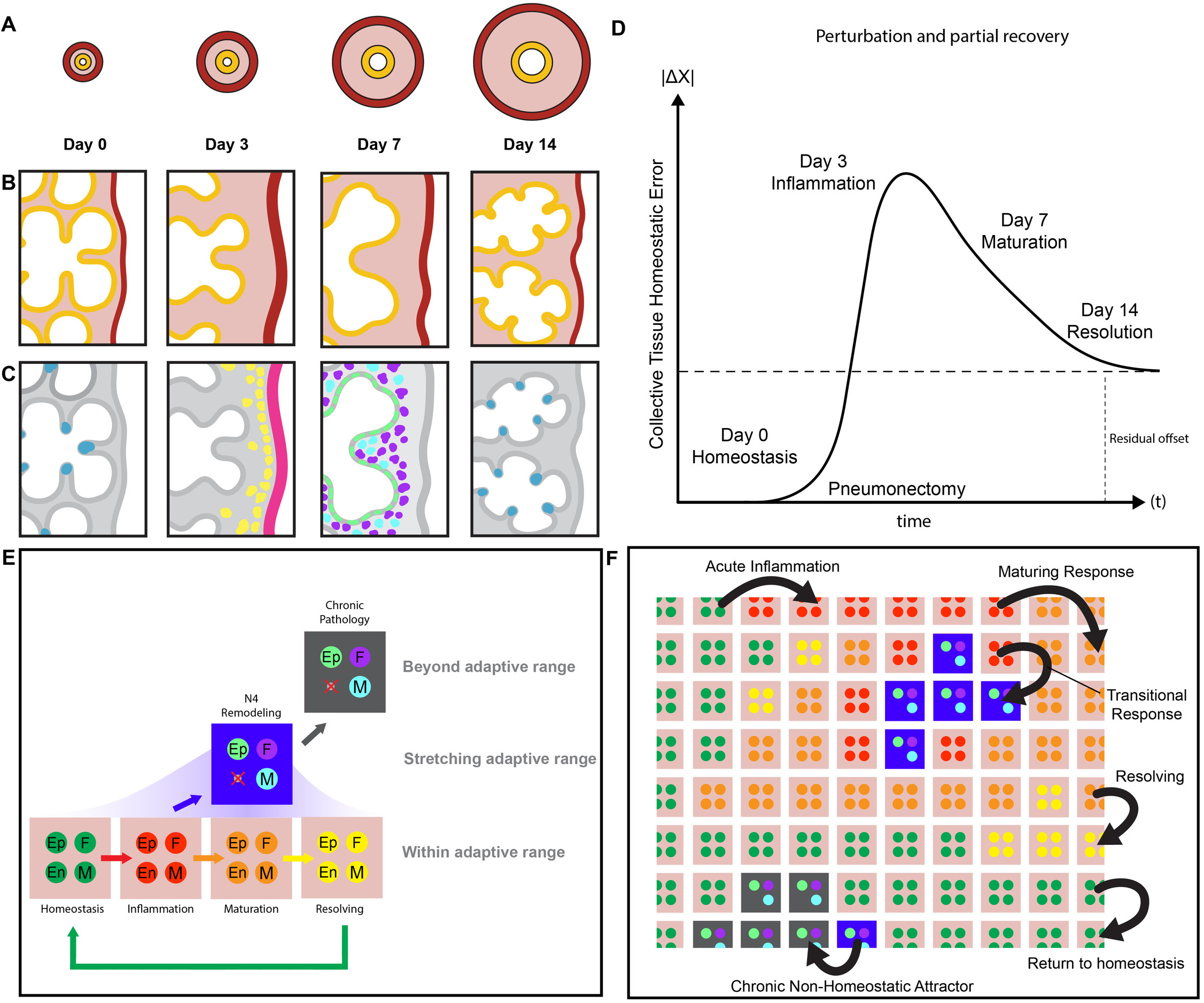
A tissue-systems model of post-pneumonectomy regeneration as traversal of a homeostatic landscape. Panels are conceptual schematics summarizing the model, not primary data. (A) Three territory simplified model of lung growth to accommodate post- pneumonectomy strain, cross section, areas not to scale in volume or surface area. (B) Schematic of distal-acinar parenchymal architecture at Day 0, 3, 7, and 14, depicting the morphologic progression from homeostatic alveoli, through inflammatory and transitional remodeling, to secondary septation. (C) Schematic cellular maps corresponding to (B), summarizing the three architectures: mesothelial activation with subpleural interstitial-macrophage accumulation at D3; the N4 transitional triad of transitional epithelium, fibroblasts, and macrophages with capillary depletion at D7; and a myofibroblast-bearing alveologenic state at D14. (D) Conceptual trajectory of collective tissue homeostatic error (|ΔX|) following pneumonectomy: a rise to peak deviation during D3 inflammation, partial recovery through D7 maturation and D14 resolution, and a residual offset reflecting the new, enlarged steady state. (E) The alveolar minimal tissue unit - epithelium (Ep), fibroblast (F), endothelium (En), macrophage (M) - traversing a homeostatic cycle *within the adaptive range* (homeostasis, inflammation, maturation, resolution, then return). Component color denotes unit state (green, homeostatic; red, inflammatory; orange, maturing; yellow, resolving). The N4 transitional state (blue) *stretches the adaptive range*, with three components adopting transitional identities while endothelium is lost (red ✕); failure to resolve permits escape *beyond the adaptive range* into a chronic fibrotic state. (F) The regenerating tissue depicted as a field of asynchronous minimal tissue units distributed across the same landscape - homeostatic (green), inflammatory (red), maturing (orange), resolving (yellow), N4-transitional (blue), and chronic non- homeostatic (dark) - connected by trajectories of acute inflammation, maturation, transitional response, resolution, return to homeostasis, and capture by a chronic non- homeostatic attractor. Because units are asynchronous in space, a single spatial snapshot samples many positions along this shared trajectory.

This framework yields several testable predictions. Resolution capacity is known to be governable by signaling alone, as when anti-TIMP1 was shown to reverse a self-reinforcing myofibroblast loop in porcine cardiac ischemia (Miyara et al., 2025). The connectomic fingerprint of the transitional state we find in lung tissues defines a candidate target set for stabilizing fibrosis-mimetic remodeling where the tissue retains regenerative competence. That fingerprint comprises gains in Spp1-Cd44, Dkk2-Lrp6, Inhba-Acvr1, and Areg-Erbb3, paired with losses in Vegfa-Kdr and Angpt1-Tek and a withdrawal of pro-resolution Bmp7 (Zou et al., 2019). These observations predict that angiogenic perturbation during PPLR should promote the transitional community, whereas exogenous vascular and maturation cues during a fibrotic challenge could accelerate resolution (Betageri et al., 2026).

By D14, the lung displays broad alveologenic maturation, with secondary septation and alveolar ringing, and secondary-crest myofibroblasts appearing alongside re-differentiated distal epithelial, endothelial, and mural populations. D14 is not simply a return to homeostasis, instead it is an active morphogenic state that carries a connectivity signature found at no other timepoint, including baseline, driven by mesenchymal-mitogen and morphogen cues (Bmp4, Fgf2, Fgf9, Hgf, Pdgfa) and Wnt potentiation (Rspo1/2).

Perhaps the most informative feature of this stage, though, is the coordinated deactivation of earlier programs. Most mechanisms perturbed earlier in the time course return within their homeostatic range by D14 (Figure 6E, green return-loop). The inflammatory and remodeling programs are actively switched off, and the tissue re-converges toward its operating point. While therapeutic attention typically falls on inhibiting the programs a tissue switches on, our data highlight a complementary property: the capacity to switch signals off is equally fundamental to the regenerative response, and the fibrotic excursion is what its failure looks like. Where quelling succeeds, the unit folds back toward resolution; where it is overwhelmed, the same transitional state settles into persistent fibrosis. Resolution in our atlas tracks the re-emergence of differentiated, gas-exchanging microvasculature at our later timepoints (Ding et al., 2011), which marks microvascular competence as a key lever for regenerative outcome.

Beyond its biological findings, this work demonstrates an approach to perturbational spatial systems biology at whole-organ scale. Responses to gross tissue perturbations are inherently complex, involving simultaneous changes in cellular composition, spatial organization, and intercellular communication. We make this complexity tractable in two complementary ways. First, formalizing each tissue section as a graph and clustering both neighborhood composition and connectivity character decomposes the response into a small number of recurring regional architectures rather than a continuum of cell states (Figure 6F). Second, layering 488 ligand–receptor connectivity profiles onto the tissue graph localizes known mechanistic literature within the ensembles where it operates, so a static snapshot can be read against functional biology rather than gene expression alone. In contrast to inference from dissociated single-cell data, which scores signaling at the cell-type level, this approach scores connectivity per cell-pair edge in real tissue space, and can be aggregated at any scale, whether cell, neighborhood, or zone, in the geometry where it occurs. The same framework extends to disease, developmental, and engineered tissues, enabling systematic cross-context comparison of multicellular signaling architecture. Where the D7 fingerprint offers targets to arrest fibrosis-mimetic remodeling, the D14 signature offers guided-morphogenesis cues for steering alveolar organoids and engineered lung tissue toward productive maturation.

Because the tissue units are asynchronous in space, a single snapshot contains units at many points along a shared trajectory. Consequently, the spatial heterogeneity we capture is not only a description of one moment but evidence of the temporal process, which Figure 6F frames our four timepoints as a field of tissue units distributed across the homeostasis, inflammation, maturation, and resolution cycle, with some on the fibrotic excursion. This is an integrative model that combines snapshot data into a coherent view of tissue adaptation over time. Within-snapshot asynchrony recovers some of the temporal resolution that four discrete timepoints would otherwise lack, but causal trajectories remain inferred rather than established. PPLR offers a spatial reference for a more general program, one that treats tissue as an agential material whose minimal tissue units sense and respond to shifts in their own setpoints, and whose guided morphogenesis we are only beginning to resolve. By mapping the same cellular and signaling primitives as they assemble into productive regeneration at one position and a fibrosis-mimetic state at another, within a single tissue, this atlas begins to specify which signals tip multicellular architecture toward regeneration and which toward pathology.

### Limitations of the study

Several limitations should be noted. First, our atlas samples only four timepoints, limiting direct reconstruction of continuous state transition trajectories at both the cellular and tissue-level. Denser temporal sampling would improve resolution of key intermediate states and circuits. Snapshot transcriptomics can support inference of dynamic and causal structure through dedicated population-dynamics and biophysical trajectory methods, none of which we apply here. Instead, temporal ordering across timepoints, combined with literature-supported connectomic localization of ligand-receptor edges, constrains interpretation of homeostatic variables that the tissue regulates and distinguishes signals that precede a response from those that follow it. Second, this study was conducted in a single species, *Rattus norvegicus*, which is less frequently used within the lung biology field. This species was chosen for its larger lung size more favorable for spatial profiling, and because it has a long history as a classical model of post-pneumonectomy lung regeneration, while also leveraging extensive prior experience with the rat model within our laboratory. Additionally, rats have a more fibrosis-prone biology (Gauldie and Kolb, 2008) and have a unique pulmonary vascular anatomy, including a more extensive intraparenchymal systemic vasculature than mice, that may influence regenerative response (Hsia et al., 2016). Future studies should compare the patterns we observe here to those in *Mus musculus*, as several of our findings are unexpected, including but not limited to the degree of parenchymal remodeling that we have captured. Cross-species studies would allow molecular perturbations and lineage traces with genetic tools that are currently not available in rat. Direct quantitative comparisons with fibrotic changes measured in *Homo sapiens* pathology would also inform candidate therapeutic targets in human disease. Third, transcript-based connectomics infers potential information transfer rather than directly measuring protein-level binding. Therefore, it is not possible to compute direct evidence of signal transduction for individual specific pathways; a known challenge in the cell-to-cell connectivity field. Future integration of spatial transcriptomics with multiplexed spatial proteomics, particularly of phosphorylated proteins, could potentially help address this limitation. Fourth, our connectivity analysis relies on a specific ground-truth database, FANTOM5, which does not include multi-subunit ligand-receptor connectivity mechanisms and thereby may allow false-positive connectivity inferences. This was a deliberate decision, as we did not want to generate false-negatives due to lack of subunit binding knowledge. However, the interpretation of the output data must be done carefully and with expert knowledge of signal transduction pathways and mediators.

Fifth, evidence of mechanical tension in the tissue as a major driver of PPLR remains indirect in our data. Future studies that independently manipulate mechanotransduction, inflammatory processes, and developmental pathways among other variables will be required to establish causal relationships among these processes. Sixth, our scRNAseq and Xenium modalities differ in cell capture: dissociation-based scRNAseq is biased toward easily dissociated alveolar populations and underrepresents cells residing within matrix-rich dense tissues and conducting structures, while Xenium captures these populations in situ. This means that some cell-type frequencies are best interpreted from the spatial data, and cross-modality comparisons should be made accounting for this bias. Seventh, under the homeostatic-perturbation lens of inflammation, inflammation and developmental circuit reactivation may be inseparable by design as cells pull from the same toolkit (Goldman and Poss, 2020). Our data cannot disentangle whether inflammatory signaling is causally required for the reactivation of developmental programs in adult tissue, or whether the two are independently co-triggered by the shared upstream stimulus of pneumonectomy and its associated mechanical and surgical perturbations. Nevertheless, the richness of the dataset, replication of our basic findings across two modalities (scRNAseq and Xenium), confirmation of several key patterns by both immunostaining and in situ hybridization, and consistent alignment with histological observations makes this dataset a robust foundation for future work.

## Resource Availability

Further information and requests for resources should be directed to and will be fulfilled by the lead contact, Micha Sam Brickman Raredon, MD PhD, available at.

## Materials availability

This study did not generate new unique reagents.

## Data and code availability

scRNA-seq and Xenium data have been deposited at GEO****** and will be made publicly available as of the date of publication. All original code has been deposited at https://github.com/RaredonLab/PPLR2026 and will be made publicly available upon the date of publication. Any additional information required to replicate the analysis of reanalyze the data reported in this paper is available from the lead contact upon request.

## Acknowledgments

The authors would like to acknowledge Ruslan Medzhitov for his longitudinal guidance and support. The authors would like to acknowledge Maor Sauler for his helpful feedback on preliminary figures. The authors would like to acknowledge Guilin Wang and William Renock at the Yale Center for Genomic Analysis for their assistance in generating single-cell and spatial transcriptomic data used in this manuscript. The authors would like to acknowledge Pavlina Baevova for her help with animal and laboratory management and Katie L. Leiby for her help with original protocol development. The authors would like to acknowledge Becky Byler and Jack Zhang for their assistance in managing controlled substances.

## Author Contributions

Conceptualization: T.O., D.P.A., M.T.G., L.E.N., and M.S.B.R.

Data Curation: T.O., D.P.A., M.T.G., S.E.E., S.M., N.W., O.P., N.Ki., K.L., and M.S.B.R.

Formal Analysis: T.O., D.P.A., M.T.G., S.E.E., S.M., N.W., O.P., N.Ki., K.L., and M.S.B.R.

Funding Acquisition: D.H.S., T.T., T.K., N.Ka., L.E.N., and M.S.B.R.

Investigation: T.O., D.P.A., M.T.G., S.E.E., S.M., N.W., N.M.S., C.D., C.H., O.P., N.Ki.,

K.L., W.D., A.M.G., T.A., R.R., H.K., V.L., Y.Y., J.C.S., A.L., A.O.Z., and M.S.B.R.

Methodology: T.O., D.P.A., S.E.E., S.M., N.M.S., C.D., C.H., J.Y., and M.S.B.R.

Project Administration: T.O., D.P.A., and M.S.B.R.

Resources: T.O., D.H.S., T.T., T.K., N.Ka., L.E.N., and M.S.B.R.

Software: D.P.A., S.E.E., N.W., J.Y., and M.S.B.R.

Supervision: T.K., N.Ka., L.E.N., and M.S.B.R.

Validation: T.O., D.P.A., M.T.G., S.E., S.M., N.W., N.M.S., C.D., O.P., N.Ki., K.L., and M.S.B.R.

Visualization: T.O., D.P.A., M.T.G., S.E.E., S.M., N.W., O.P., N.Ki., K.L., and M.S.B.R.

Writing – Original Draft: T.O., D.P.A., M.T.G., S.E.E., and M.S.B.R.

Writing – Review & Editing: D.P.A., M.T.G., S.E.E., O.P., N.Ki., K.L., W.D., and M.S.B.R.

## Declaration of generative AI and AI-assisted technologies in the writing process

During the preparation of this work the author(s) used Anthropic’s Claude (Claude Opus 4.8, with earlier Claude Opus and Sonnet versions in sessions prior to May 2026) in order to edit manuscript prose for clarity, rhythm, and stylistic consistency, draft new paragraphs from author-specified content briefs, and reorganize manuscript structure under author direction. After using this tool, the author(s) reviewed and edited the content and take full responsibility for the content of the published article.

## STAR★Methods

## Methods

### In vivo

#### Animal husbandry

All rats were wild-type Sprague Dawley rats (*Rattus Norvegicus*) obtained from Charles River (8-10 weeks old, 200-350 g) and bred and maintained at the animal center of Yale University. All animal procedures (pneumonectomy survival surgery and terminal surgery) were approved by the Yale University Institutional Animal Care and Use Committee (IACUC protocol #2025-11190) and performed in accordance with the NIH Guidelines for the Care and Use of Laboratory Animals. Male and female animals were used. The study was not specifically powered to detect sex-based effects, which may limit the ability to identify biological differences and should be considered when interpreting generalizability.

#### Pneumonectomy

Sprague Dawley rats of both sexes (8-10 weeks) were anesthetized with ketamine (75 mg/kg) and xylazine (5 mg/kg), orally intubated with a 16G catheter, and maintained using Isoflurane via mechanical ventilation throughout the procedure. All procedures were performed under aseptic conditions, with continuous pulse oximetry monitoring. The left chest was shaved, disinfected with ethanol and iodine, and immobilized in the right lateral position. Thoracotomy was performed via left intercostal caudal to the left scapula (at 4th-5th intercostal space). All membranous structures attached to the left lobe were dissected, leaving only the pulmonary hilum attached, and the pulmonary artery/vein and the left main bronchus were then collectively ligated using 7-0 polypropylene thread. The entire left lobe was excised and discarded. The chest was closed, anesthesia withdrawn and post-surgical analgesia provided with Buprenorphine Sustained Release. NSAIDs were withheld to avoid systemic anti-inflammatory effects. Animals were monitored daily until scheduled euthanasia.

#### Rat lung dissociation

At days 3, 7, and 14 post-pneumonectomy, animals were anesthetized with Ketamine (75 mg/kg) and Xylazine (5 mg/kg) and injected IP with sodium heparin (400 U/kg). The abdomen was opened in sterile fashion and the xiphoid process lifted to reveal the underside of the diaphragm. The diaphragm was punctured and retracted to reveal the chest cavity. The rib cage was lifted, and an incision was made axially to reveal the trachea. The trachea was separated from the esophagus, opened using fine scissors, cannulated with a barbed fitting, and secured with 4-0 silk. The right ventricle was opened with scissors, and the pulmonary artery (PA) cannulated with an 18-gauge feeding needle and secured with 4-0 monofilament polypropylene. An incision was made in the left ventricle, and the lungs were perfused via the PA with 100 mL of PBS containing 100 U/mL sodium heparin and 0.01 mg/mL sodium nitroprusside with concurrent airway ventilation. The heart-lung block was then removed from the chest to be perfused with enzyme.

Tissue dissociation was performed using DMEM containing 1mg/mL Collagenase/Dispase (Roche, Cat No. 11097113001), 3 U/mL Elastase (Worthington, Cat No. LS002292), and 20 U/mL DNase (Qiagen, Cat No. 79254) pre-heated to 37°C. Enzyme was perfused through the vasculature and then instilled into the trachea 5-10x. Care was taken to fully recruit and inflate the alveolar regions of the lung and to have the tissue fully saturated with enzyme. Following inflation and/or perfusion with enzyme, non-lung tissue was removed, including the heart and extralobular airways. The remaining tissue and effluent were placed on a rocker at 37°C and 60 rpm for 20 minutes. At this point the lung tissue was pushed through a fine wire sieve using a weighing spatula until only dense connective tissue remained. The strainer was rinsed with 20 mL ice-cold DMEM containing 10% FBS, 1% penicillin/streptomycin (Gibco, Cat No. 15140-122), 1% amphotericin (HyClone, Cat No. SH30071.03), and 0.1% gentamicin (Gemini Bio, Cat No. 50-753-3028). The resulting tissue solution was spun down for 5 min at 300 x g, the supernatant discarded, and the pellet resuspended in ACK lysing buffer (Gibco, Cat No. A1049201) at a 1:1 ratio with pellet volume for 120 seconds at room temperature. Following dilution and re-centrifugation, the pellet was resuspended in 0.01% BSA in PBS and filtered sequentially through a 70 µm filter (Falcon, Cat No. 352350) and twice through a 40 µm filter (Falcon, Cat No. 352340). The resulting cell suspension was counted, assessed for viability using 0.4% Trypan Blue solution (Gibco, Cat No. 15250061), and serially diluted to the desired concentration for single cell library preparation.

#### Single-cell library preparation and sequencing

Single-cell RNA sequencing libraries were generated using the Chromium Single Cell 3’ v2 and v3 assays (10x Genomics) and an in-house Drop-seq platform. Samples were diluted targeting a recovery of 10,000 cells per lane. Libraries were sequenced on the HiSeq4000 platform (Illumina) to a target depth of 50,000 reads per cell.

#### Rat lung fixation

Surgically isolated rat lungs were cleared of blood, inflated with approximately 5 mL of air (half-inflation), and perfused intravascularly via the PA with room-temperature 4% paraformaldehyde (PFA) delivered at 30 cmH2O via gravity with recirculation. Perfusion continued for a minimum of 3 hours. Tissue was subsequently immersed in 4% PFA for 12 hours at room temperature, then transferred to 70% ethanol for storage and downstream processing.

#### TMA creation

A 3x8 tissue microarray (TMA) was constructed by Yale Pathology Tissue Services. We selected 24 unique 2 mm biopsy punches (n=12 animals; 4 females and 8 males), with two samples per animal, one subpleural and one adventitial. These tissue punches were assembled into a single paraffin TMA, enabling the profiling of all samples within a single Xenium slide.

#### Immunochemistry on paraffin sections

Samples of native rat lung were fixed in formalin for 3 hours, then paraffin-embedded and sectioned at Yale Pathology Tissue Services (YPTS). Hematoxylin and eosin (H&E) staining of sections was carried out by YPTS or in-house. Formalin-fixed paraffin-embedded (FFPE) sections were deparaffinized by heating at 65°C for 30 minutes, followed by a graded xylene and ethanol series (100%, 95%, 70%). Antigen retrieval was performed in citric acid buffer (0.01 M citric acid, 0.05% Tween-20, pH 6.0) at 75°C for 20 minutes. After cooling, sections were rinsed in PBS and permeabilized for 15 min in permeabilization solution (0.2% Triton X-100 (Invitrogen, Cat No. HFH10) in PBS). Primary antibodies (see list of antibodies used in Supplement 1: Key Resources Table) were applied overnight at 4°C in blocking buffer. Primary antibodies were diluted in blocking buffer as follows: rabbit anti-Cd163 (ab182422)(1:100), rabbit anti-YAP (D8H1X)(Cell Signaling Technologies, Cat No. 14074)(1:200), rabbit anti-Cxcl6 (Invitrogen, Cat No. PA5-115276)(1:50). Following PBS washes, sections were incubated in Alexa Fluor conjugated secondary antibodies at 1:500 dilution for 1 hour at room temperature, counterstained with DAPI for 1 min, and mounted with PVA-DABCO (Sigma Aldrich, Cat No. 10981).

#### mRNA FISH

A total of six formalin-fixed, paraffin-embedded rat lung tissue sections (4% PFA buffered in 0.1 M sodium phosphate buffer) were analyzed, including accessory lobes from Day 0 and Day 3 post-pneumonectomy. For each timepoint, slides were assigned to positive control probes (Ubc, Ppib, and Polr2a), negative control probes (dapB), or target probes.

Slides were incubated at 60 °C for 1 hour, deparaffinized with Xylene and 100% ethanol, and air dried. RNAscope assays were performed according to the manufacturer’s protocol. Briefly, sections were treated sequentially with hydrogen peroxide, target retrieval reagent (100°C), and Protease Plus (40°C). Control slides were incubated with species-specific positive or negative control probes, while experimental slides were hybridized with probe mixtures (C2 and C3 probes diluted into C1 probe solutions). Hybridization was performed for 2 hours at 40°C in a HybEZ oven. Slides were then washed and stored overnight in 5X SSC buffer at room temperature. Following overnight incubation with 5X SSC buffer, signal amplification was performed using AMP1, AMP2, and AMP3 reagents at 40 °C with intermediate washing steps.

HRP-based signal development was performed sequentially for each channel, with fluorophore labeling using TSA-based dyes for Wnt4 (C1), Angptl4 (C2), and Bmp4 (C3). Between each channel, HRP blocking steps were applied according to the manufacturer’s instructions. Slides were counterstained with DAPI, mounted using Prolong Gold Antifade and stored overnight at 2-8 °C. Slides were imaged using a Nikon Eclipse Ji fluorescence microscope.

### In Silico

#### Single-cell data extraction and filtration

Raw sequencing data were aligned and extracted using STARsolo and the Ensembl Rnor6.0 release. The top 30,000 cell barcodes per sample, ranked by nUMI count, were then imported into R for further processing.

#### Single-cell data cleaning, clustering and annotation

Single-cell RNA-seq data (∼90,000 cells, 4,000 median UMIs/cell) were generated using Drop-seq and 10x Genomics platforms (v2 and v3.1). Each sample was conservatively filtered independently to remove low-quality cells and technical artifacts. Cleaned datasets were then integrated and subjected to iterative clustering, with removal of low-information clusters and putative multiplets. Cells were subsequently assigned to major classes based on canonical marker expression (epithelial, endothelial, lymphoid, myeloid, or mesenchymal), and each class was independently subclustered and annotated, yielding 51 distinct cell types.

#### Statistical analysis of single-cell data

To identify genes exhibiting robust and sex-conserved regulation across the post-pneumonectomy time course, differential expression analyses were performed in a stratified manner. The dataset was first partitioned by sex and subsequently by cell type annotation, yielding 68 independent R objects (34 cell types × 2 sexes). Within each subset, differential expression across time points was computed using the Wilcoxon rank-sum test implemented in Seurat (*FindAllMarkers*), applying minimal filtering thresholds (min.cells.feature = 3, min.cells.group = 3, min.pct = 0, logfc.threshold = 0, return.thresh = 1, only.pos = FALSE) to retain a comprehensive set of candidate genes.

Two large data frames were assembled, each capturing significance testing results from each sex. Then, each data fram was annotated with additional metadata describing unique row identifiers (gene-CellType-Timepoint combinations) and added columns designating if a given row was found to be up or down regulated in each sex. Sex-specific data frames were then merged to generate a consensus dataset preserving all observations across conditions. For each feature, mean log fold changes were computed across sexes, and p-values were combined using Fisher’s method. Statistical significance required concordant directionality of differential expression between sexes and a combined adjusted p-value < 0.05.

#### Gene panel design for spatial transcriptomics

A targeted gene panel was curated to enable spatial mapping of cellular identity and intercellular communication. To ensure accurate cell type and state annotation in spatial data, 107 genes were first included as markers of lineage and cell states. An additional 373 genes encoding ligands and receptors were included based on differential expression and ligand–receptor inference analyses, collectively capturing 488 unique ligand–receptor mechanisms annotated in the FANTOM5 database and predicted to be perturbed during PPLR. Gene selection was guided by combined statistical significance, magnitude of differential expression, and consistency of perturbation across sexes.

#### Xenium 10X data extraction and filtration

Raw Xenium transcript data were exported from YCGA in Apache Parquet format and were converted to gzip-compressed CSV format for importing. Data were imported into R using LoadXenium() with a molecule quality threshold of QV ≥ 20, inclusion of cell centroids and molecule coordinates, and cell segmentations provided by the platform. Basic quality control was performed by visualizing transcript and feature counts per cell. Since multiple TMA cores were arrayed across the single Xenium capture area, individual TMA regions were manually cropped based on spatial coordinates from Xenium Explorer software. Cell segmentations were verified post-cropping. Preprocessed data were normalized, variable features identified and scaled in Seurat. PCA was performed, and 37 PCs were selected based on elbow plot inspection and PC heatmap evaluation for UMAP embedding, neighborhood graph construction, and Louvain clustering (res = 0.3). Cell centroid coordinates and TMA identifiers were appended to the metadata, and major cell classes (epithelial, endothelial, mesenchymal, and immune) were initially validated using canonical markers (Epcam, Cdh5, Col1a1, Ptprc). Of the 330,540 total cells detected across the slide, 330,317 were successfully assigned to a named TMA region; the remaining 223 cells, which fell outside all defined crop boundaries, were excluded from subsequent analyses.

#### Xenium 10X data clustering and annotation

Xenium spatial transcriptomic data (∼330,000 cells, mean 250 counts per cell) were generated using the 10x Genomics Xenium In Situ platform. Following image-based segmentation and transcript assignment, low-quality cells and segmentation artifacts were removed through iterative quality control. Cells were initially clustered into main cell classes based on canonical marker expression (epithelial, endothelial, immune, and mesenchymal), and ambiguous low- information or putative doublet populations were manually reviewed during iterative re-clustering. Each cell class was then independently subclustered and annotated using canonical marker genes, spatial localization patterns, and reference comparisons to matched single-cell RNA-seq datasets from the same experimental system, yielding 59 distinct cell states.

#### Connectomics analysis

Ligand–receptor mediated intercellular communication was inferred using the NICHES framework, applied independently to both the single-cell RNA-seq dataset and the Xenium spatial transcriptomics dataset.

#### Tissue graph construction and Neighborhood Composition Quantification

We experimented with multiple tissue graph-construction approaches and empirically determined that a local-radius based approach (ε-graph) was preferable to a nearest-neighbors approach (kNN). The resulting graphs have the desirable property of maintaining edge density proportional to cellular density while avoiding mapping edges between cells far enough that diffusion-mediated ligand-receptor communication would be unreasonable. These graphs were then used to compute a local neighborhood assay for each cell, counting the number of adjacent cells classified as each cell type. Finally, hyperedge data was created for each edge, characterizing each cell-to-cell dyad within high-dimensional ligand-receptor connectivity space. We organized these data into three primary assays from which to perform more complex downstream analysis: Xenium (single-cell spatial transcriptomics, rows = genes, columns = cells), Cell-to-Cell (single-edge spatial connectomics, rows = mechanisms, columns = edges), and Neighborhood (single-cell spatial communities, rows = annotated cell states, columns = cells). These three data layers allowed us to connect local cellular state (Xenium) to local cellular connectivity (CTC) and local cellular community (Neighborhood).

Spatial neighbor graphs were constructed independently for each TMA to avoid artifactual connections between distinct tissue sections. For each TMA, pairwise Euclidean distances between all cell centroids (x, y coordinates) were computed, and edges were drawn between all cell pairs within a fixed radius of 25 µm. This threshold approximates one to two cell diameters in the lung parenchyma, capturing biologically meaningful direct cell-cell contacts. A radius-based definition was preferred over a k-nearest neighbor approach, as the latter imposes uniform neighbor counts across cells regardless of local density, an inadequate assumption for PPLR pucks where cellular density significantly varies across regions. Spatial graphs were exported as weighted edge lists via the *igraph* package, with edge weights reflecting inter-centroid Euclidean distances.

Cell type neighborhood composition was then quantified by projecting metadata annotations onto each graph. For every cell, neighboring cell types were tallied to produce both a raw count vector and a fractional composition vector normalized by total neighbor count. Results were computed per TMA, concatenated, and stored as objects for downstream analysis. All analyses used custom R functions built on the Seurat, dplyr, tidyr, and igraph packages.

## QUANTIFICATION AND STATISTICAL ANALYSIS

### Units of analysis and level of replication

The spatial dataset is hierarchically structured: transcripts are nested within cells, cells within TMA pucks, and pucks within animals, with each animal contributing exactly two pucks (one subpleural and one adventitial). Because cells within a puck and pucks within an animal are not independent observations, treating individual cells as replicates would constitute pseudoreplication and would produce anticonservative p-values inflated by several orders of magnitude relative to the true biological sample size. Accordingly, all inferential statistics reported for tissue-level composition and abundance use the puck as the observational unit and the animal as the unit of biological replication, with animal identity entered as a random intercept. Cell-level and edge-level quantities (per-cell neighbor densities, per-edge ligand-receptor scores) are reported descriptively, or are aggregated to the puck or animal level before any hypothesis test is performed. Where a per-cell statistic is presented directly, it is explicitly labeled as descriptive and no p-value is attached to it.

### Software and reproducibility

All analyses were performed in R. Single-cell and spatial data structures were handled with Seurat and SeuratObject; graph construction used igraph; data manipulation used dplyr and tidyr. Mixed-effects models were fit with lme4 and glmmTMB; model diagnostics used DHARMa and performance; estimated marginal means and contrasts used emmeans; smooth trends were fit with mgcv via ggplot2. Differential expression used Seurat, muscat, and iDESC. Random seeds were set globally prior to any resampling procedure so that bootstrap and permutation results are exactly reproducible. The complete analysis environment is captured in a package lockfile accompanying the code repository, and all analysis scripts, custom functions, and shared configuration (including all palettes and thresholds) are deposited with the manuscript.

### Neighborhood assay construction and unsupervised neighborhood clustering

The Neighborhood assay described above (rows = annotated cell states, columns = cells, values = count of neighbors of each type within the 25 um radial graph) was deliberately not library-size normalized. Each column is already an interpretable count of physically adjacent cells, and normalizing would have discarded the biologically meaningful variation in local cellular density that distinguishes dense fibrotic foci from sparse alveolar parenchyma. Raw counts were therefore carried directly into the data layer and z-scaled across features prior to dimensionality reduction. Twelve features corresponding to low-information populations and putative segmentation doublets were excluded from the feature set used for embedding, on the grounds that these labels do not represent discrete, reliably identifiable cell states and would introduce segmentation artifact into the neighborhood definition; they remain present in the object and in all descriptive composition summaries. Principal component analysis was run on the retained features, and 29 components were selected on the basis of elbow plot inspection and component heatmap evaluation. A shared nearest-neighbor graph was constructed on these components and clustered by the Louvain algorithm at resolution 0.2. Clustering resolution was chosen to produce neighborhood classes that were spatially contiguous and interpretable on the tissue; the resulting cluster identities are used throughout as categorical labels, and no statistical claim depends on the specific resolution parameter.

### Histologic territory assignment

Coarse histologic territories (Pleural, Parenchymal, Conducting) were assigned by nearest-landmark proximity. For every cell, the Euclidean distance to the nearest cell of each user-defined landmark cell type was computed, and the cell was assigned to the territory corresponding to its closest landmark. Distances were computed strictly within TMA puck, since inter-puck distances on the array reflect the geometry of the tissue microarray rather than tissue anatomy and are biologically meaningless. Cells with no landmark within the search domain were left unassigned rather than forced into the nearest category, and unassigned cells are reported explicitly rather than silently dropped.

### Mixed-effects modeling of neighborhood abundance across the time course

To test whether the abundance of a given spatial neighborhood changes across the post-pneumonectomy time course, cell-level neighborhood labels were aggregated to a per-puck summary table in which each row records the number of cells assigned to the neighborhood of interest and the number assigned to any other neighborhood. Animal identity was derived by stripping the regional suffix from the puck identifier, and data integrity was verified by confirming that every puck mapped to a valid animal and that every animal contributed exactly two pucks, one per region.

Neighborhood abundance was modeled as a binomial proportion using a generalized linear mixed-effects model of the form cbind(neighborhood cells, other cells) ∼ Timepoint + Region + Sex + (1 | Animal), fit by maximum likelihood with a logit link. The binomial response formulation weights each puck by its total cell count, so that pucks contributing more cells carry proportionally more information, while the logit link constrains fitted proportions to the unit interval. The random intercept for animal accounts for the paired subpleural and adventitial samples drawn from each animal and is the term that prevents pseudoreplication. Timepoint was treated as an ordered factor with Day 0 as the reference level. Models were fit with the bobyqa optimizer and an increased iteration limit to ensure stable convergence on this small, clustered design; every fit was checked for convergence warnings and for singularity of the random-effects covariance.

The necessity of the random intercept was verified rather than assumed. The intraclass correlation coefficient was computed from the fitted binomial model as the ratio of between-animal variance to total variance on the logit scale, using the standard logistic-distribution residual variance of pi-squared over three, and confirmed that a non-trivial fraction of variance is attributable to animal identity.

Because sex was unbalanced with respect to timepoint by design, the sex-by-timepoint cross-tabulation was inspected before interpretation, and sex is retained in the model as a covariate to adjust for composition rather than as a term for which we make any inferential claim.

### Model selection, dispersion, and diagnostics

Individual model terms were tested by likelihood ratio test comparing the full model against a reduced model omitting a single term, rather than by relying on Wald z-statistics, which are unreliable at small numbers of clusters. A Timepoint-by-Region interaction was fit and formally compared against the additive model by likelihood ratio test. A random slope for region within animal was additionally attempted to allow the regional effect to vary by animal; this specification was singular given the number of animals available and was therefore not retained, and the simpler random-intercept structure was carried forward.

Model adequacy was assessed using simulation-based scaled residuals (DHARMa), which compare the observed residuals against residuals simulated from the fitted model. The formal dispersion test indicated overdispersion relative to the binomial expectation, meaning the observed puck-to-puck variability exceeded that permitted by a binomial likelihood. Rather than report the anticonservative binomial p-values, primary inference was moved to a beta-binomial mixed model (glmmTMB, logit link) with the identical fixed and random effects structure, which admits an additional dispersion parameter and yields appropriately widened intervals. The interaction test was repeated in the beta-binomial specification before the final model structure was fixed. The binomial fit is retained only as supporting evidence for the intraclass correlation calculation.

### Effect sizes, contrasts, and multiplicity

Effects are reported as risk differences on the response scale (absolute differences in predicted proportion, in percentage points) rather than as odds ratios, because odds ratios are difficult to interpret for outcomes of this magnitude and are not the quantity of biological interest. Estimated marginal means were computed from the final model with emmeans and regridded to the response scale before contrasts were formed. Because the sampling distribution of a risk difference derived from a mixed model with few clusters is not reliably normal, confidence intervals and p-values for all contrasts were obtained by parametric bootstrap with 2,000 replicates. Families of pairwise contrasts were corrected for multiple comparisons using the Holm procedure, which controls the family-wise error rate without assuming independence among contrasts. Where an interaction was retained, simple effects (timepoint within region, and region within timepoint) are reported instead of main effects.

We note explicitly that this design is small: with the number of animals available, the bootstrap Holm-adjusted contrasts on neighborhood abundance did not reach conventional significance, notwithstanding a consistent rise-peak-decline trajectory with a maximum near Day 7. We therefore describe these temporal patterns as trends supported by consistent effect direction and by the convergent spatial analyses below, and we do not claim statistical significance for them. Exact estimates, confidence intervals, and adjusted p-values for every contrast are reported in the corresponding figure legends and supplementary tables.

### Decomposition of abundance into nucleation and expansion

A change in the mean abundance of a neighborhood across time can arise either because the neighborhood appears in more samples (nucleation) or because it occupies a larger fraction of the samples in which it is already present (expansion). Because these have distinct biological interpretations and are conflated in a single proportion model, the outcome was additionally modeled as a two-part (hurdle) decomposition. Nucleation was modeled as a logistic mixed-effects regression on the binary indicator of whether the neighborhood was present at all in a given puck, with the same fixed effects and animal random intercept. Expansion was modeled as the binomial proportion model described above, restricted to pucks in which the neighborhood was present. Separation in the presence-absence cross-tabulation was checked before fitting the nucleation model, since complete or quasi-complete separation renders logistic coefficients unbounded and their standard errors uninterpretable.

### Spatial covariance between vascular and transitional-triad density

To test the prediction that local vascular supply and local transitional-cell burden are inversely related within regenerating parenchyma, two per-cell quantities were computed for every parenchymal cell from the precomputed 25 um radial edge list: local vascular density, defined as the fraction of that cell’s graph neighbors annotated as gCap or aCap; and local transitional-triad density, defined as the fraction of neighbors annotated as Transitional-Epithelium, Transitional-Fibroblasts, or Transitional-Macrophage. Both are computed from the existing graph, so no additional radius or bandwidth parameter was introduced and no further tuning was performed.

The association between these two quantities was quantified by Spearman rank correlation, which makes no assumption of linearity or bivariate normality and is robust to the zero-inflated, right-skewed distribution of local density fractions. Confidence intervals were obtained by resampling cells with replacement (500 bootstrap replicates), taking the 2.5th and 97.5th percentiles of the resulting distribution of rho.

### Stratified correlation analysis and the effective sample size

A correlation computed by pooling all parenchymal cells at a timepoint has a nominal n in the tens of thousands but an effective n set by the number of independent tissue samples, and a pooled estimate can in principle be produced entirely by between-puck differences rather than by within-tissue co-variation. To address this directly, the Spearman correlation was recomputed at three nested stratifications rather than reported only as a pooled value. First, per puck, yielding one rho per puck per timepoint, each estimated from thousands of cells within a single tissue section, so that co-variation is demonstrated within samples rather than across them.

Second, per puck type, separating subpleural from adventitial pucks at each timepoint. Third, per animal, pairing the subpleural and adventitial pucks contributed by the same animal. Each stratified estimate carries its own bootstrap confidence interval, and the distribution of per-puck rho values is reported directly so that the consistency of the effect across independent biological samples can be assessed by inspection rather than inferred from a single pooled statistic.

The per-puck-type stratification additionally serves as a control against a surgical-artifact explanation. Adventitial pucks are not subject to the pleural and thoracotomy geometry that could plausibly generate a spurious vascular-to-transitional gradient; recovery of the same negative association in adventitial pucks therefore argues against that interpretation.

### Distance-to-pleura gradient analysis

For each parenchymal cell, the Euclidean distance to the nearest pleural mesothelial cell was computed strictly within the same TMA puck, for the reasons given above. Relationships between pleural distance and each regenerative metric were visualized in two complementary ways that do not depend on the same assumptions. Continuous trends were fit with a generalized additive model using a thin-plate regression spline with the basis dimension deliberately restricted to four, which permits a single monotone or single-peaked trend while preventing the smoother from chasing noise at the sparse, long-distance tail of the distribution. In parallel, pleural distance was discretized into five quantile-defined bands per timepoint and the distribution of each metric within each band was displayed as a violin plot. Quantile banding rather than fixed-width binning was used so that every band contains a comparable number of cells, avoiding the unstable estimates that fixed-width bins produce in the sparsely populated tail. These smooths and bands are descriptive summaries of spatial organization; formal inference on gradient differences across timepoints is not claimed from them.

### Differential expression in the spatial dataset

Differential expression across the spatial time course was assessed with methods that model sample-level structure explicitly, rather than with cell-level rank tests, which treat every cell as an independent replicate and are known to be severely anticonservative in multi-sample single-cell designs. Mixed-model differential state analysis was performed per cell type using muscat, contrasting each timepoint against all others with a variance-partitioning linear mixed model (the dream framework) that carries a random effect for sample. In parallel, pseudobulk profiles were generated by summing raw counts within each cell type and sample, and differential expression was tested on these pseudobulk profiles under explicit design and contrast matrices, both for pairwise timepoint comparisons and for one-timepoint-versus-rest comparisons. Pseudobulk aggregation reduces the effective sample size to the number of biological samples and is the more conservative of the two approaches; agreement between the mixed-model and pseudobulk results was used as the criterion for reporting a gene as differentially expressed. As an additional check on genes of interest, two-group comparisons were repeated with iDESC, which explicitly models subject-level variation in single-cell count data, using the TMA puck as the subject identifier and applying minimum cell-per-subject and minimum-subject thresholds to exclude comparisons that are underpowered at the subject level.

### Aggregation and standardization of ligand-receptor mechanism scores

Ligand-receptor mechanism (LRM) scores are defined per edge and are not directly comparable across mechanisms, because mechanisms differ by orders of magnitude in the expression level of their constituent genes. Before any aggregation, each mechanism’s edge scores were therefore standardized to zero mean and unit variance across the entire edge population, placing all mechanisms on a common scale of standard deviations from the dataset-wide typical value. Standardized scores were then averaged within the stratum of interest (histologic territory, neighborhood class, and timepoint), retaining only edges for which both the sending and the receiving cell belong to that stratum, so that reported signatures reflect signaling internal to the compartment rather than signaling across its boundary. The resulting signatures are descriptive summaries of relative signaling composition and are presented as such.

Because the assignment of cells to a putatively pathological neighborhood is itself an analytical choice that could drive the result, the compartment-level signaling analysis was run under three prespecified alternative treatments of that neighborhood: excluding its cells from all territories, including them, and comparing them directly against territory-matched cells outside it, each computed both pooled across timepoints and stratified by timepoint. Reporting all three treatments rather than a single chosen one makes the dependence of the conclusion on that analytical decision explicit and auditable.

### Permutation testing for spatial association

Where the question was whether a categorical spatial label is associated with a timepoint or with a feature more than expected by chance, significance was assessed by label permutation rather than by a parametric test, since the null distribution of such spatial association statistics is not analytically tractable and the underlying observations are spatially autocorrelated. The observed test statistic was compared against an empirical null distribution generated by randomly permuting the label of interest 10,000 times while holding all other structure fixed, and a two-sided empirical p-value was computed as the proportion of permuted statistics at least as extreme as the observed statistic in either direction.

### Statistical limitations

Several limitations bound the strength of the inferential claims made here and are stated explicitly. The number of animals is small, which limits power for tissue-level contrasts; where contrasts did not reach significance we report effect direction and magnitude with intervals rather than asserting a null result, since absence of significance at this sample size is not evidence of absence of effect. All spatial samples were profiled on a single Xenium capture area, which eliminates run-to-run batch effects by construction but also means that no run-level batch term can be estimated and any hypothetical slide-level artifact would be perfectly confounded with the experiment as a whole. Sex is included as a model covariate but the study was not powered to detect sex-based effects, and no sex-specific claim is made from the spatial data. Cell-level and edge-level statistics involve very large nominal n but a much smaller effective n, and are reported descriptively or aggregated before testing for that reason. Finally, neighborhood classes and cell type annotations are the product of unsupervised clustering followed by expert curation; downstream statistics are conditional on those assignments and should be interpreted as such.

## KEY RESOURCES TABLE

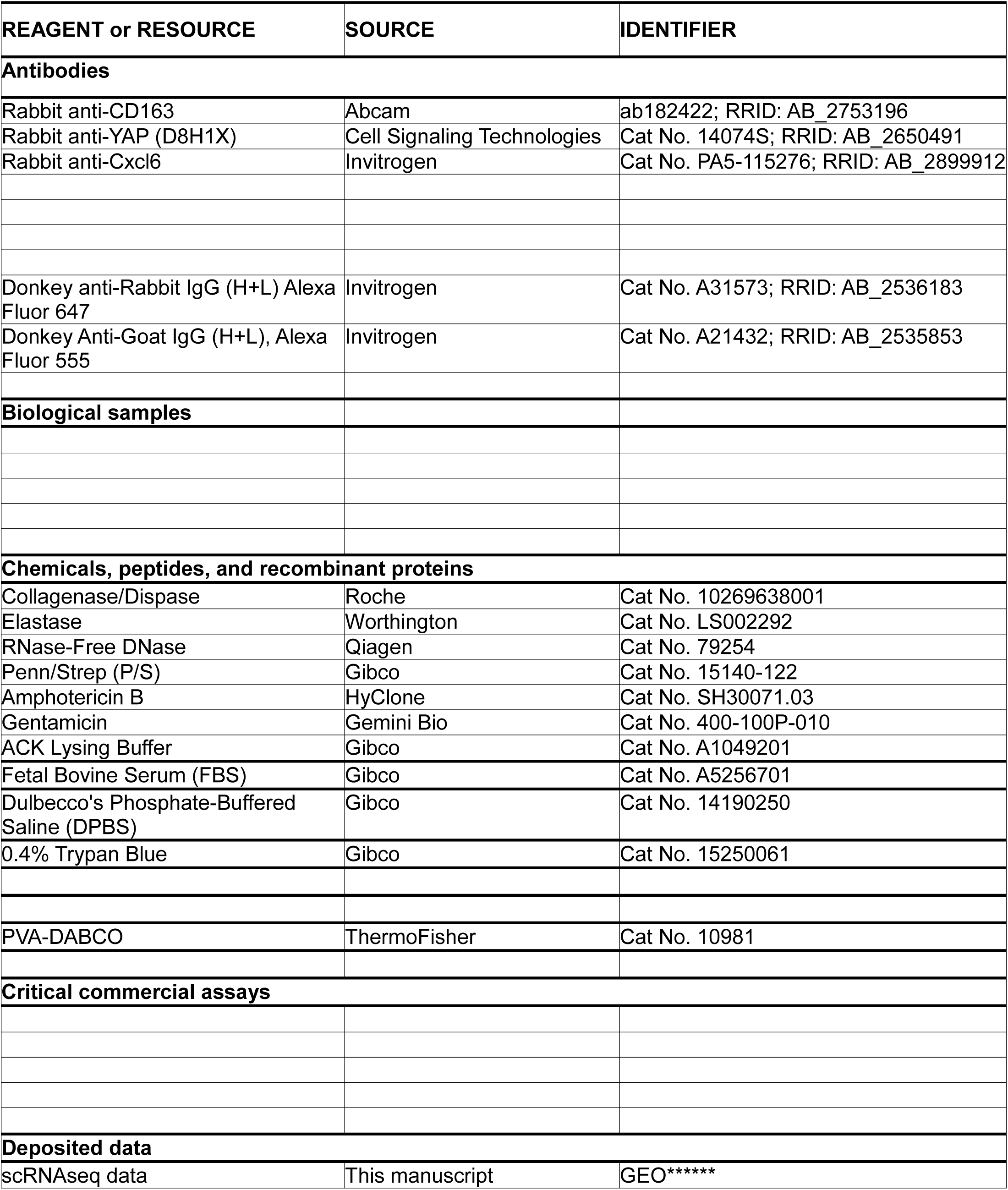
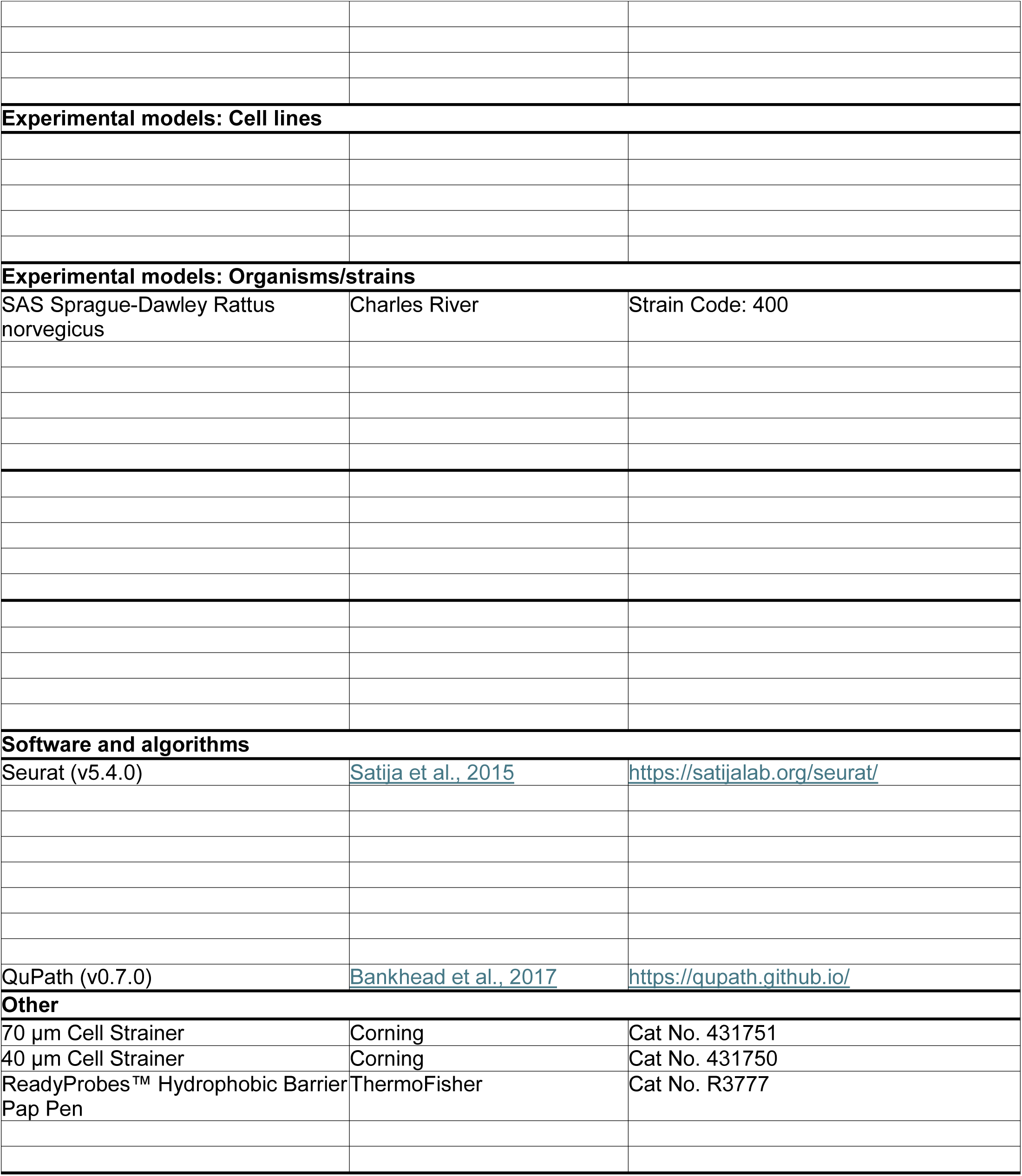

---

**Figure S1.**
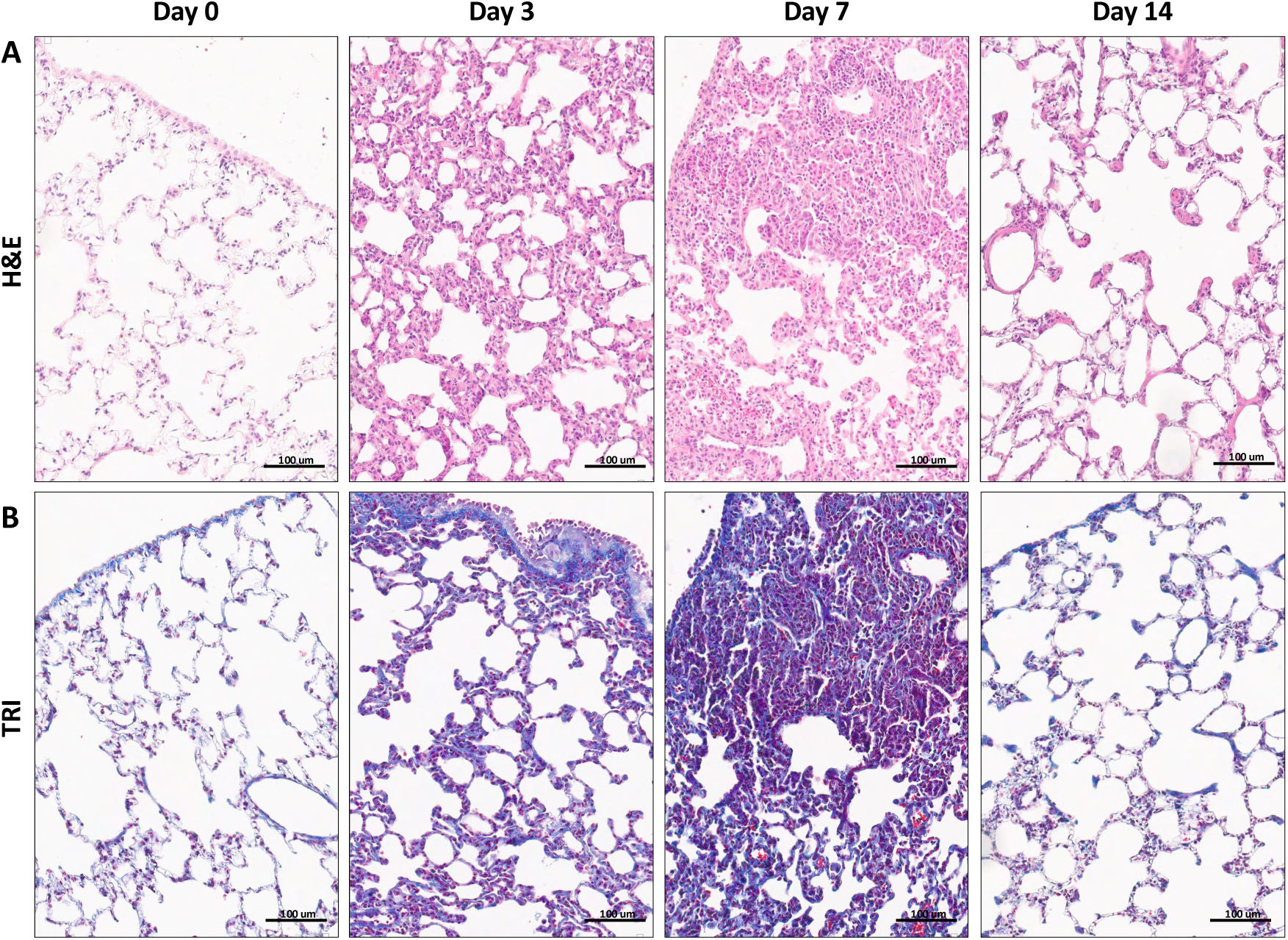
Histology across the post-pneumonectomy time course. Representative histological sections of rat lung tissue at Day 0 (homeostasis), Day 3, Day 7, and Day 14 post-pneumonectomy, stained with (A) hematoxylin and eosin (H&E) and (B) Masson’s trichrome (TRI). At Day 0, alveolar architecture is intact, with thin gas-exchanging septa and open airspaces. By Day 3, parenchyma shows widespread inflammatory changes, including septal thickening and mononuclear infiltrate. By Day 7, select parenchymal regions display dense cellular consolidation with marked loss of alveolar architecture, accompanied by increased trichrome staining indicating matrix and cellular density. By Day 14, alveolar architecture is largely restored, with thin septa and open airspaces resembling baseline homeostasis. Scale bars: 100 μm.

**Supplemental Figure 2.**
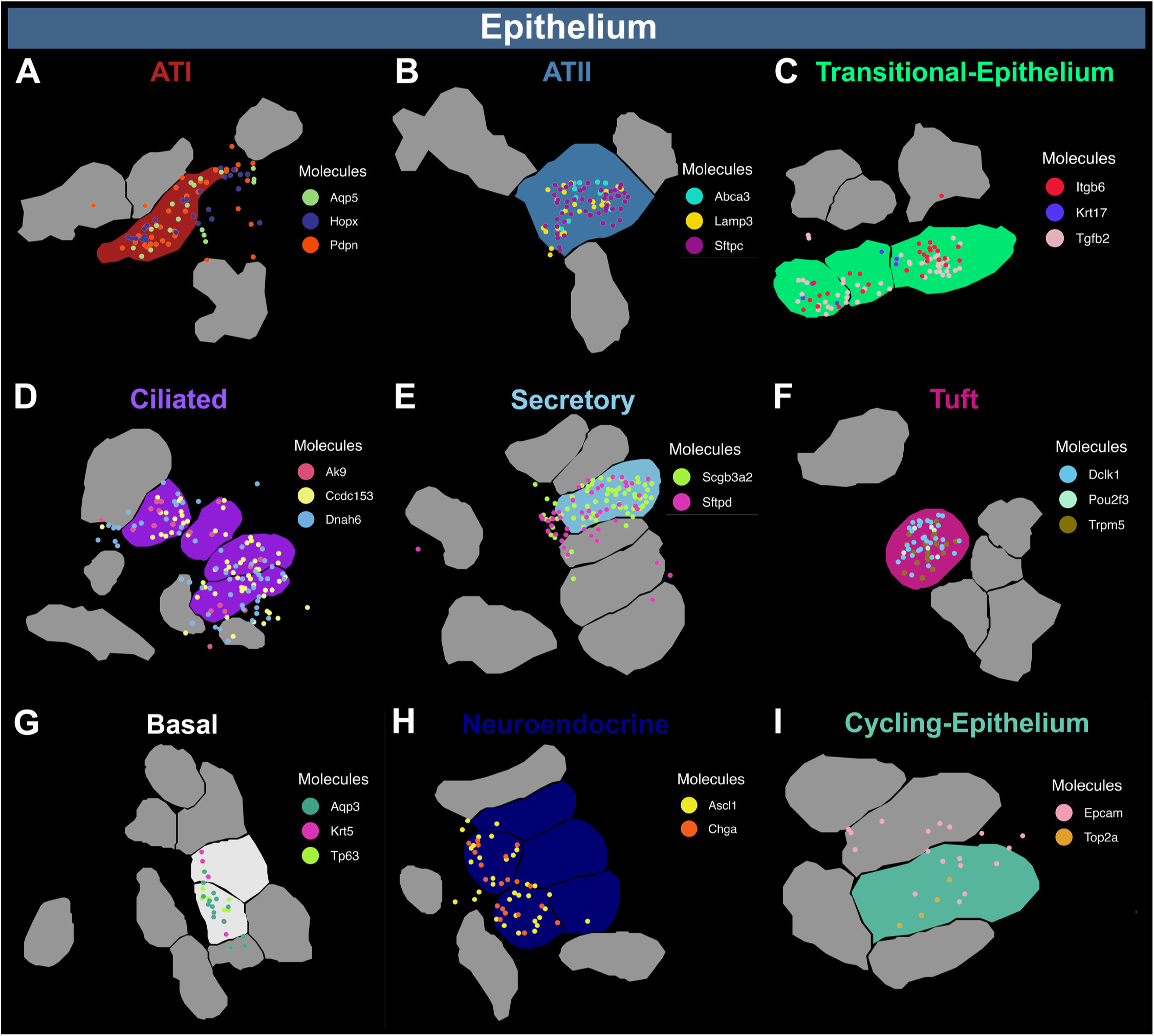
Epithelial cell populations identified in the Xenium spatial atlas through expression of canonical lineage-associated genes during PPLR. The following cell types are shown: (A) ATI, (B) ATII, (C), Transitional-Epithelium, (D) Ciliated, (E) Secretory, (F) Tuft, (G) Basal, (H) Neuroendocrine, and (I) Cycling-Epithelium. Colored segmented cells indicate the annotated epithelial population of interest, while neighboring cells are shown in gray. Overlaid transcript coordinates demonstrate expression of representative canonical marker genes commonly used to define each epithelial lineage or transcriptional state.

**Supplemental Figure 3.**
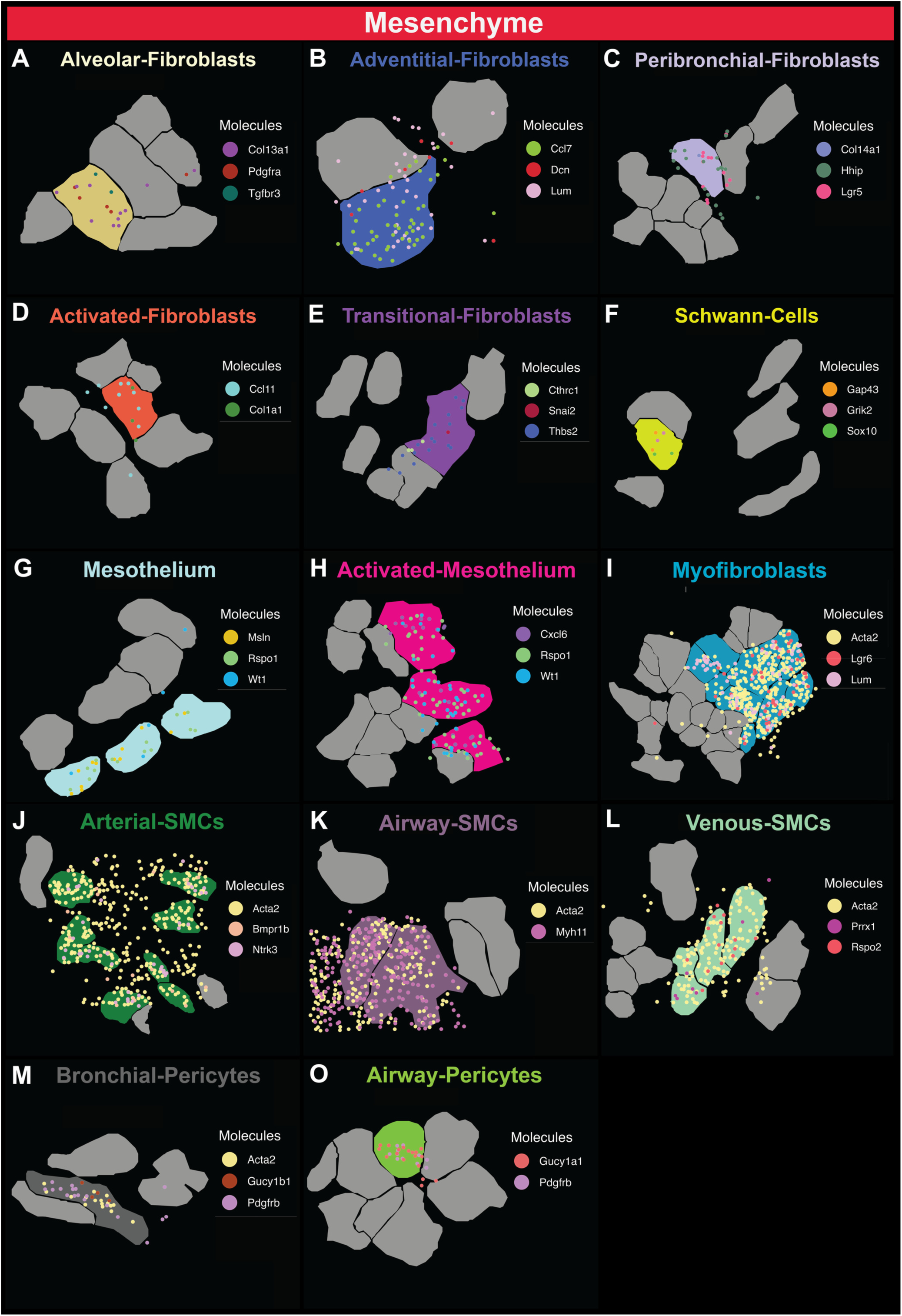
Mesenchymal cell populations identified in the Xenium spatial atlas through expression of canonical lineage-associated genes during PPLR. The following cell types are shown: (A) Alveolar-Fibroblasts, (B) Adventitial-Fibroblasts, (C) Peribronchial-Fibroblasts, (D) Activated-Fibroblasts, (E) Transitional-Fibroblasts, (F) Schwann-Cells, (G) Mesothelium, (H) Activated-Mesothelium, (I) Myofibroblasts, (J) Arterial-SMCS, (K) Airway-SMCs, (L) Venous-SMCs, (M) Bronchial-Pericytes, and (N) Airway-Pericytes. Colored segmented cells indicate the annotated mesenchymal population of interest, while neighboring cells are shown in gray. Overlaid transcript coordinates demonstrate expression of representative canonical marker genes commonly used to define each mesenchymal subtype.

**Supplemental Figure 4.**
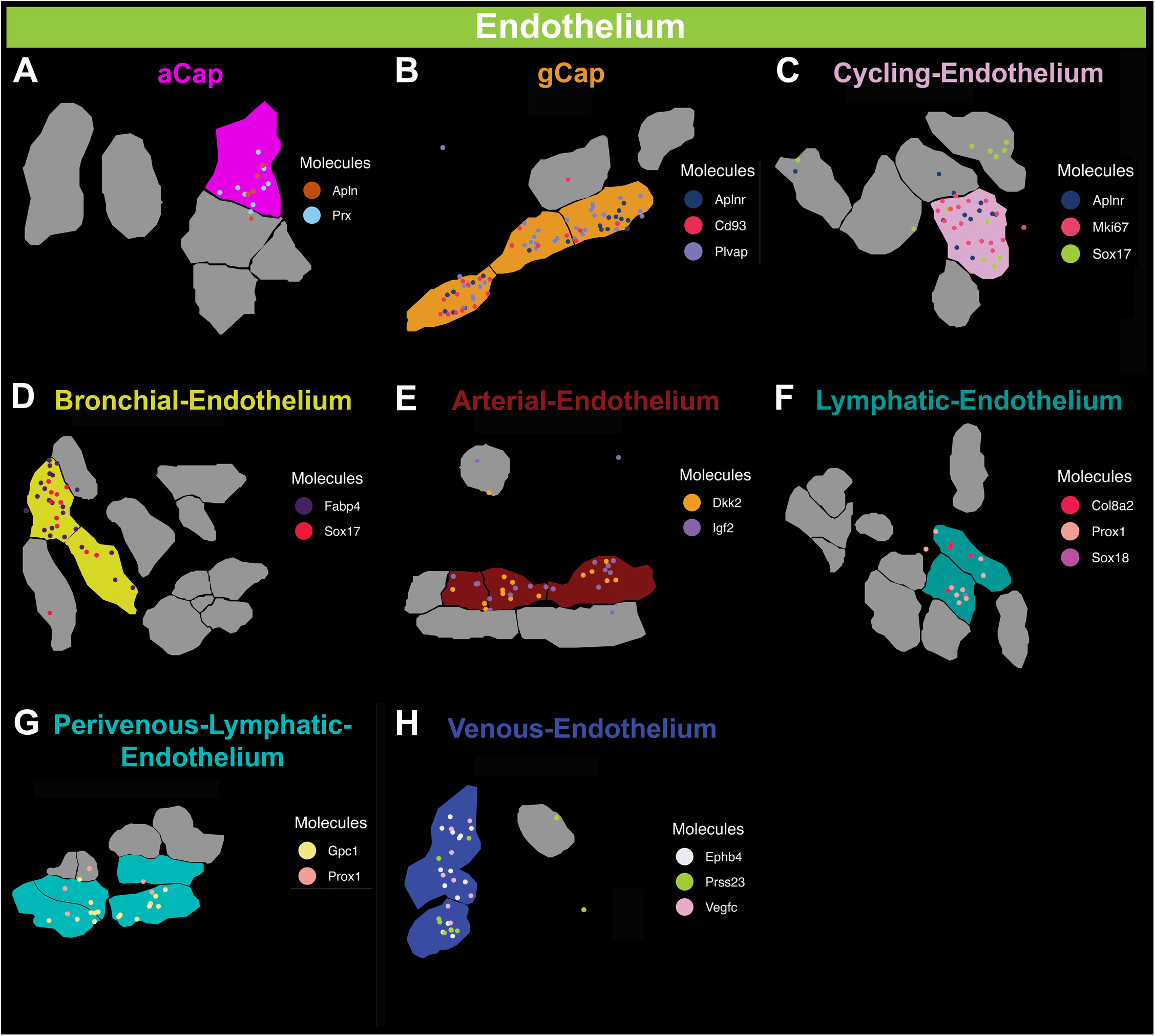
Endothelial cell populations identified in the Xenium spatial atlas through expression of canonical lineage-associated genes during PPLR. The following cell types are shown: (A) aCap, (B) gCap, (C) Cycling-Endothelium, (D) Bronchial-Endothelium, (E) Arterial-Endothelium, (F) Lymphatic-Endothelium, (G) Perivenous-Lymphatic-Endothelium, and (H) Venous-Endothelium. Colored segmented cells indicate the annotated endothelial population of interest, while neighboring cells are shown in gray. Overlaid transcript coordinates demonstrate expression of representative canonical marker genes commonly used to define each endothelial subtype.

**Supplemental Figure 5.**
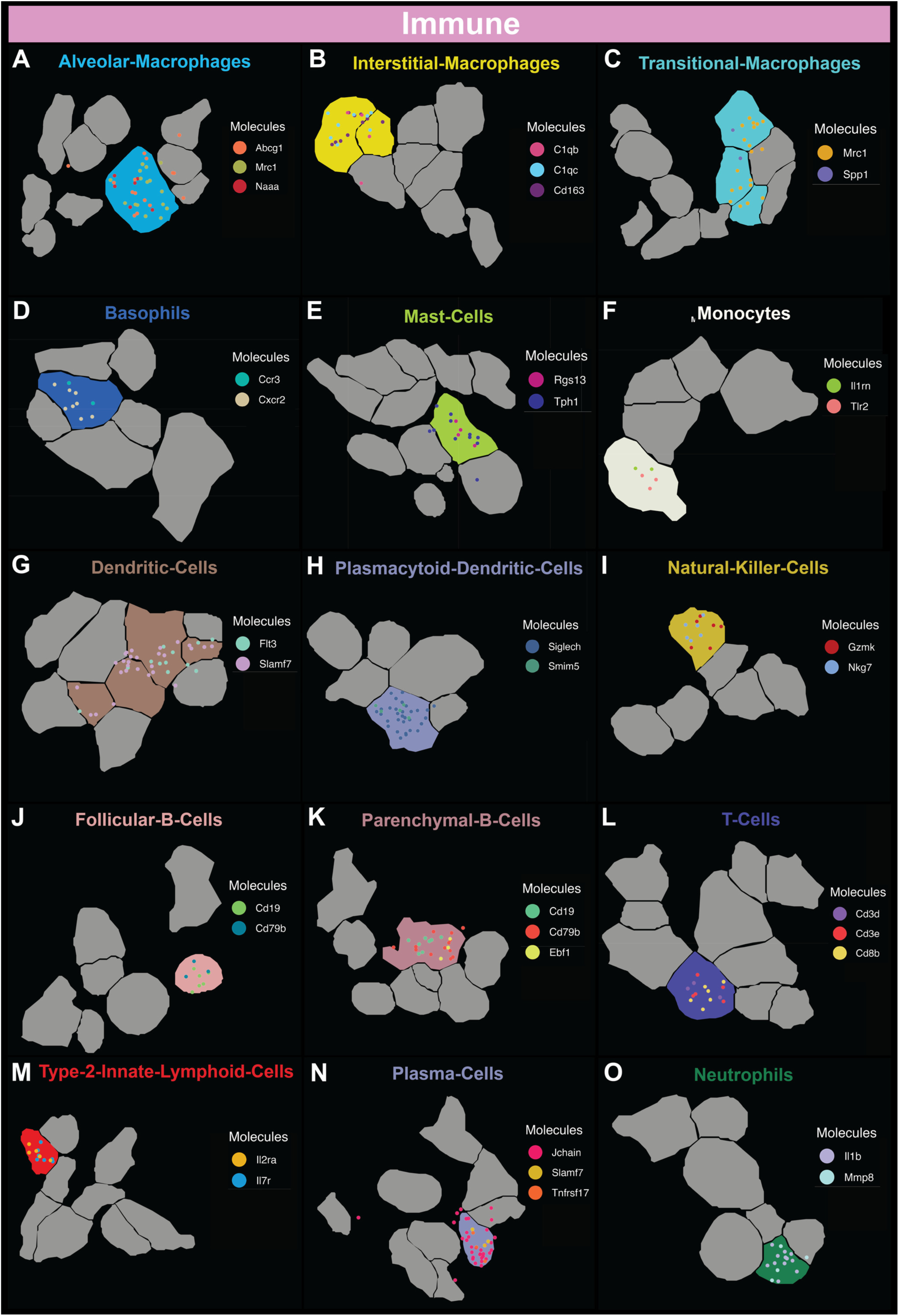
Immune cell populations identified in the Xenium spatial atlas through expression of canonical lineage-associated genes during PPLR. The following cell types are shown: (A) Alveolar-Macrophages, (B) Interstitial-Macrophages, (C) Transitional-Macrophages, (D) Basophils, (E) Mast-Cells, (F) Monocytes, (G) Dendritic- Cells, (H) Plasmacytoid-Dendritic-Cells, (I) Natural-Killer-Cells, (J) Follicular-B-Cells (K) Parenchymal-B-Cells, (L) T-Cells, (M) Type-2-Innate-Lymphoid-Cells, (N) Plasma-Cells, and (O) Neutrophils. Colored segmented cells indicate the annotated immune population of interest, while neighboring cells are shown in gray. Overlaid transcript coordinates demonstrate expression of representative canonical marker genes commonly used to define each immune subtype.

**Supplemental Figure 6.**
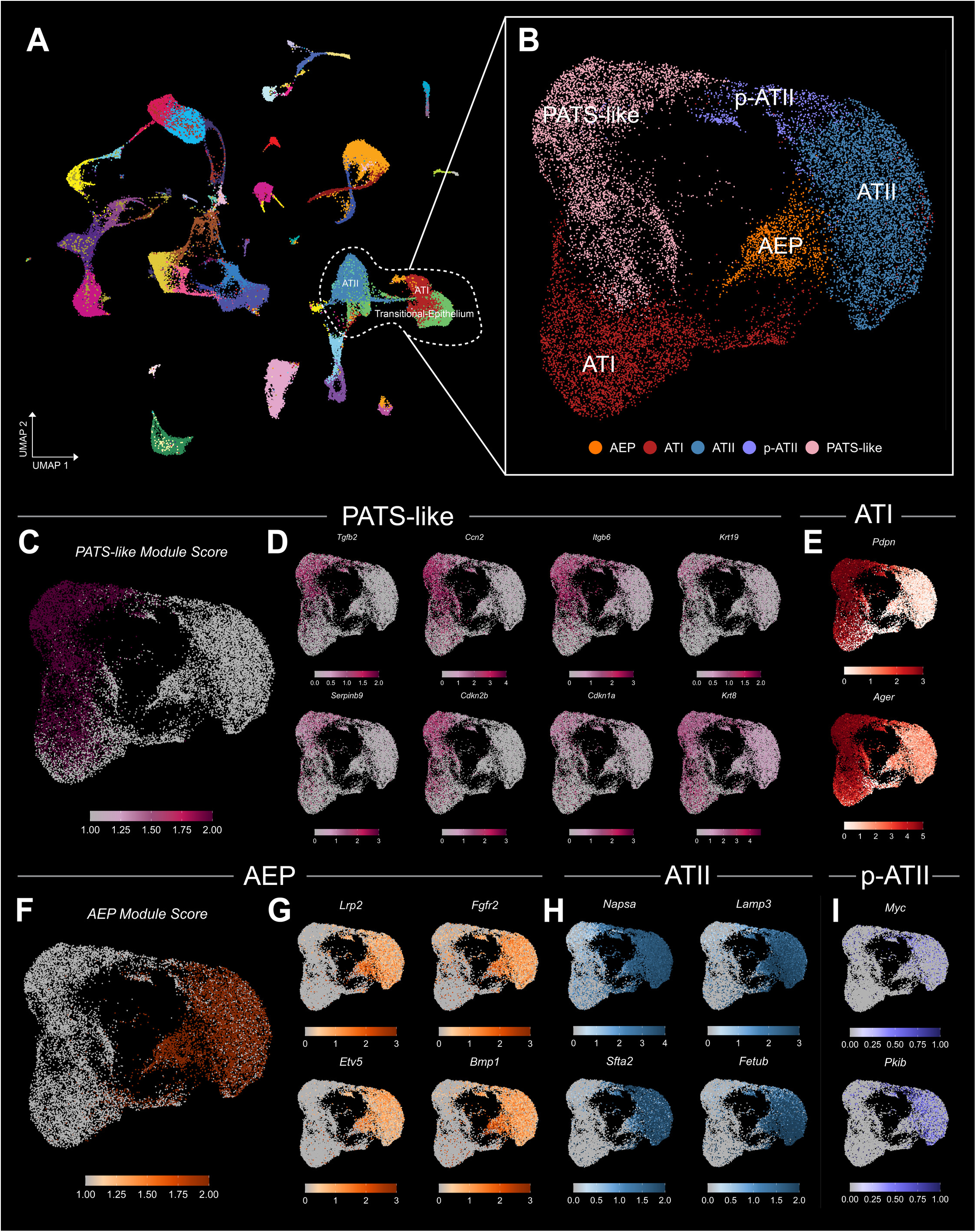
Sub-clustering analysis of epithelial populations in scRNAseq dataset to identify transcriptionally distinct transitional epithelial states analogous to previously described regenerative-associated epithelial programs. A) UMAP visualization of the integrated scRNA-seq atlas with the distal epithelial cells used for subset analysis outlined. B) UMAP embedding of subsetted ATI, ATII, and Transitional- Epithelium populations following re-integration and re-clustering analysis, identifying ATI, ATII, proliferating-ATII (p-ATII), alveolar epithelial progenitor-like (AEP-like), and pre-alveolar type-1 transitional-like states (PATS-like) based on shared transcriptional features with previously reported epithelial programs. C) Module score visualization for the PATS-associated program. D) Representative feature plots showing marker expression patterns supporting annotation of PATS-like cells. E) Representative feature plots showing canonical markers for ATI cells. F) Module score visualization for the AEP-associated program. G-I) Representative feature plots showing marker expression patterns supporting annotation of AEPs (G), ATII cells (H), and p-ATII cells (I). Gene lists used for AEP and PATS-like module scores can be found in Supplemental Spreadsheet 1.

**Figure S7.**
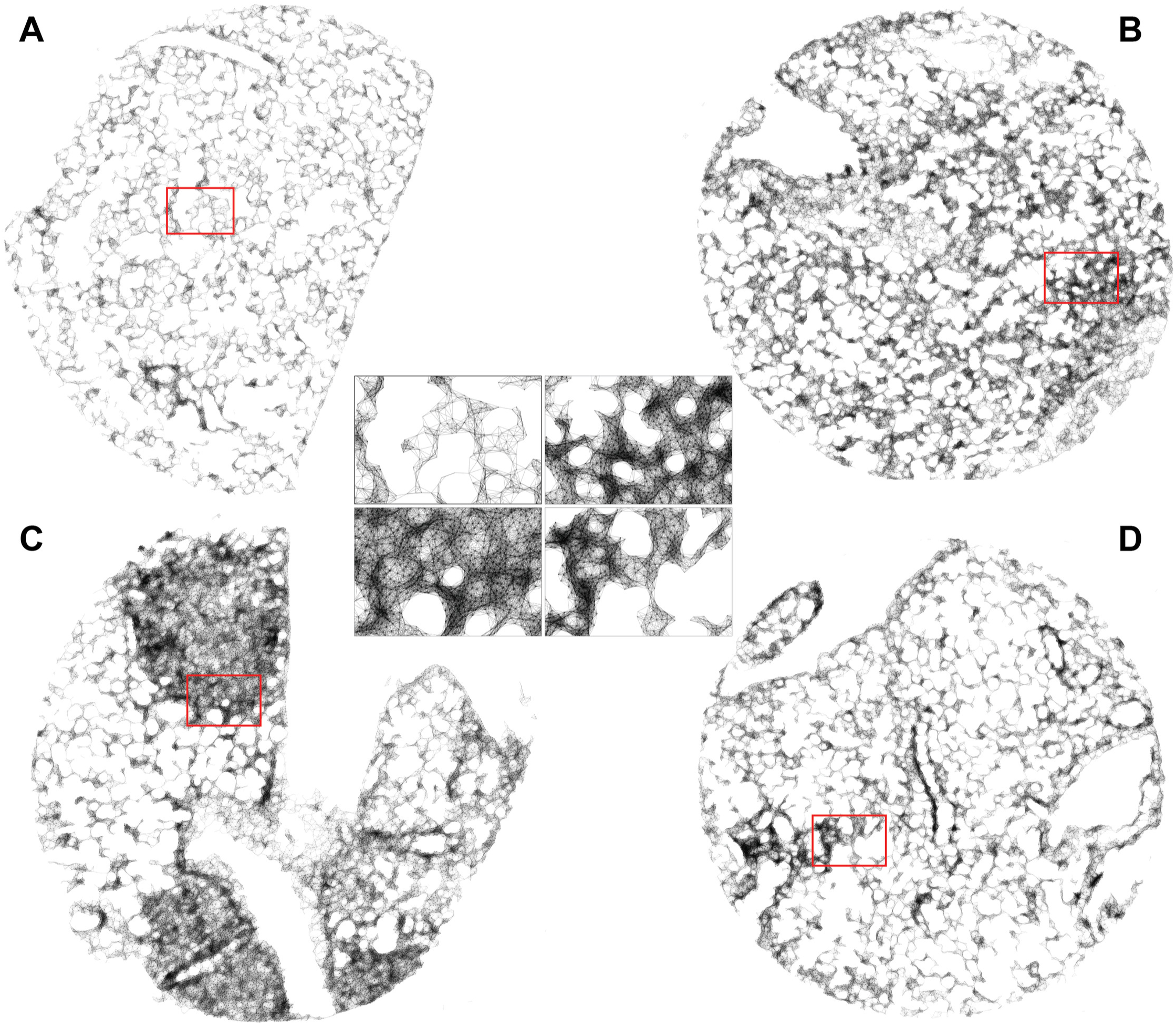
Tissue graphs across the regeneration timecourse. **(A–D)** Representative tissue graphs from Day 0, 3, 7, and 14, respectively. Insets show the boxed region from each panel at higher magnification to illustrate graph structure. Graphs encode only edge information between cells, not cell identity, constructed using a local radius-based approach: two cells are connected by an edge if they lie within 25 µm of one another, a distance consistent with the diffusion range of ligands in lung tissue. The resulting graph approximates tissue architecture while preserving alveolar, airway, and vessel lumens. Because edges reflect cellular colocalization rather than a fixed number of nearest neighbors, graph density also serves as a high-yield proxy for local tissue density. The regions shown here were selected to illustrate the changes in local tissue density observed over the course of lung regeneration.

**Figure S8.**
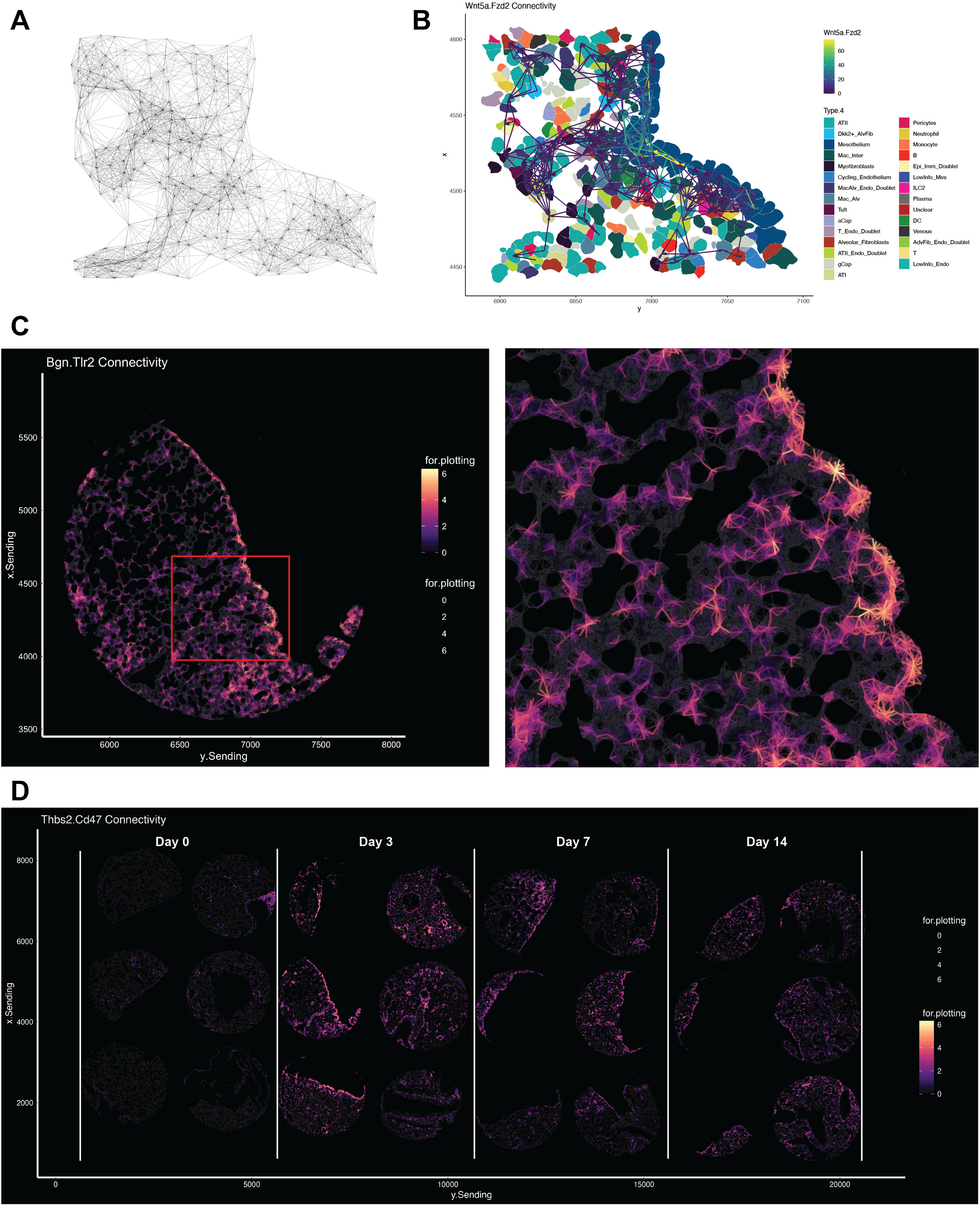
Ligand–receptor connectivity overlays on tissue graphs. **(A)** Base tissue graph for a selected region of tissue (Day 3 pleural region, chosen for known mesothelial activation and local subpleural interstitial macrophage enrichment). **(B)** A single connectivity mechanism (*Wnt5a*-*Fzd2*) overlaid on this graph, colored by intensity, computed as the product of ligand and receptor expression in the sending and receiving cell, respectively. **(C)** A different connectivity mechanism (*Bgn*-*Tlr2*) shown at whole-puck scale, illustrating that these connectivity graphs allow precise spatial localization of regions relatively enriched or depleted for a given signal. Inset at right emphasizes graph structure; gray edges denote cell-cell edges that were mapped but carry zero connectivity for the selected mechanism. **(D)** *Thbs2*-*Cd47* connectivity across the full tissue microarray, capturing change over time across all six replicates per timepoint; these data show that pneumonectomy induces widespread regenerative circuitry changes with readily observable spatiotemporal dynamics and heterogeneous enrichment. This dataset contains equivalent connectivity layers for 488 unique ligand-receptor mechanisms (LRMs), and statistical analysis of these layers underlies many of the findings presented in this manuscript.

**Figure S9.**
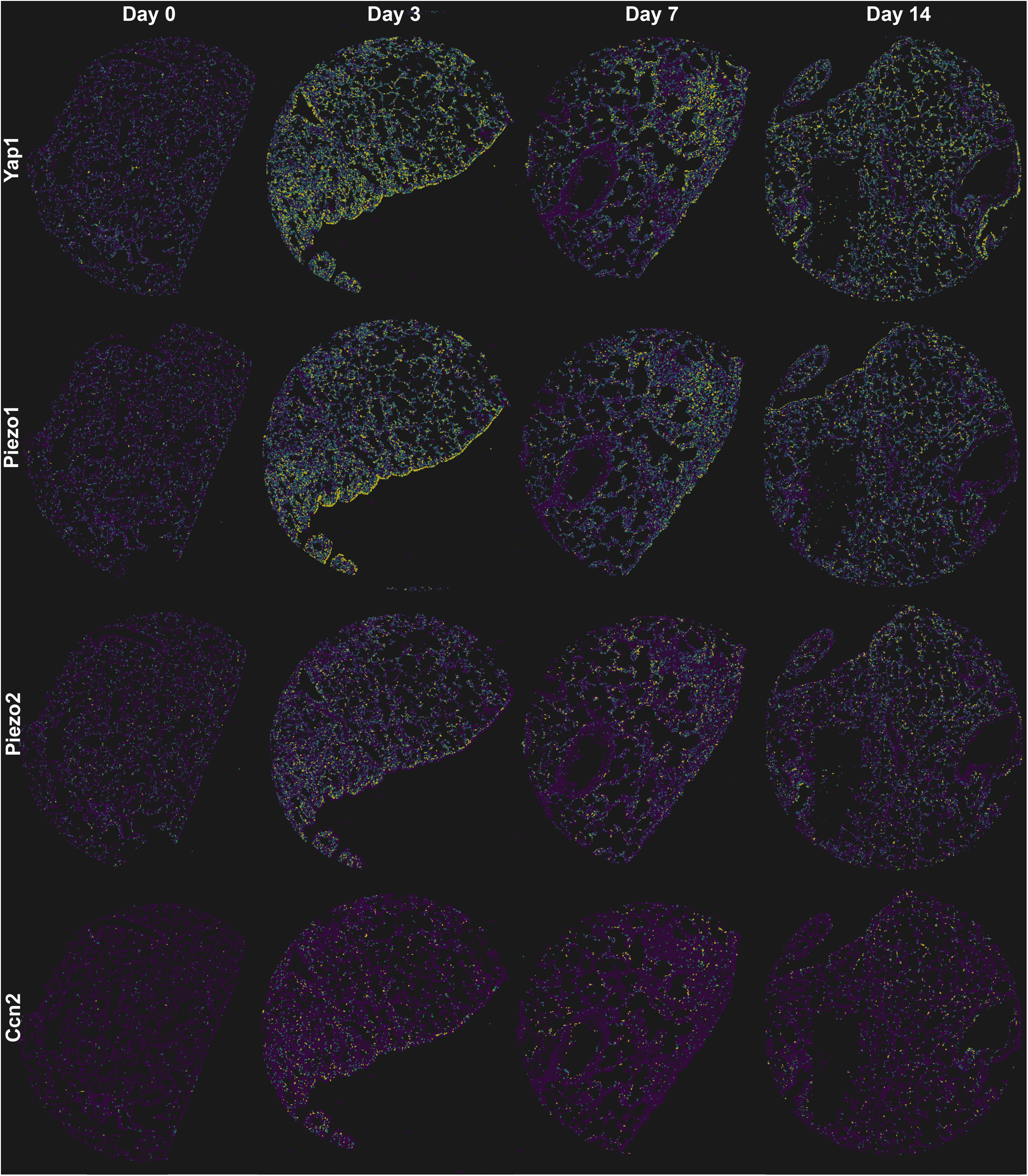
Hippo pathway activation over time. Representative tissue pucks from Day 0, 3, 7, and 14 (one per timepoint), with cells colored by gene expression. Four genes are shown: *Yap1*, *Piezo1*, *Piezo2*, and *Ccn2* (*Ctgf*), all established markers of cellular mechanotransduction in the lung. Expression of all four genes rises sharply by Day 3, remains elevated during the fibromimetic remodeling observed at Day 7, and shows sparse, localized downregulation by Day 14, coincident with alveolarization and alveolar maturation.

**Figure S10.**
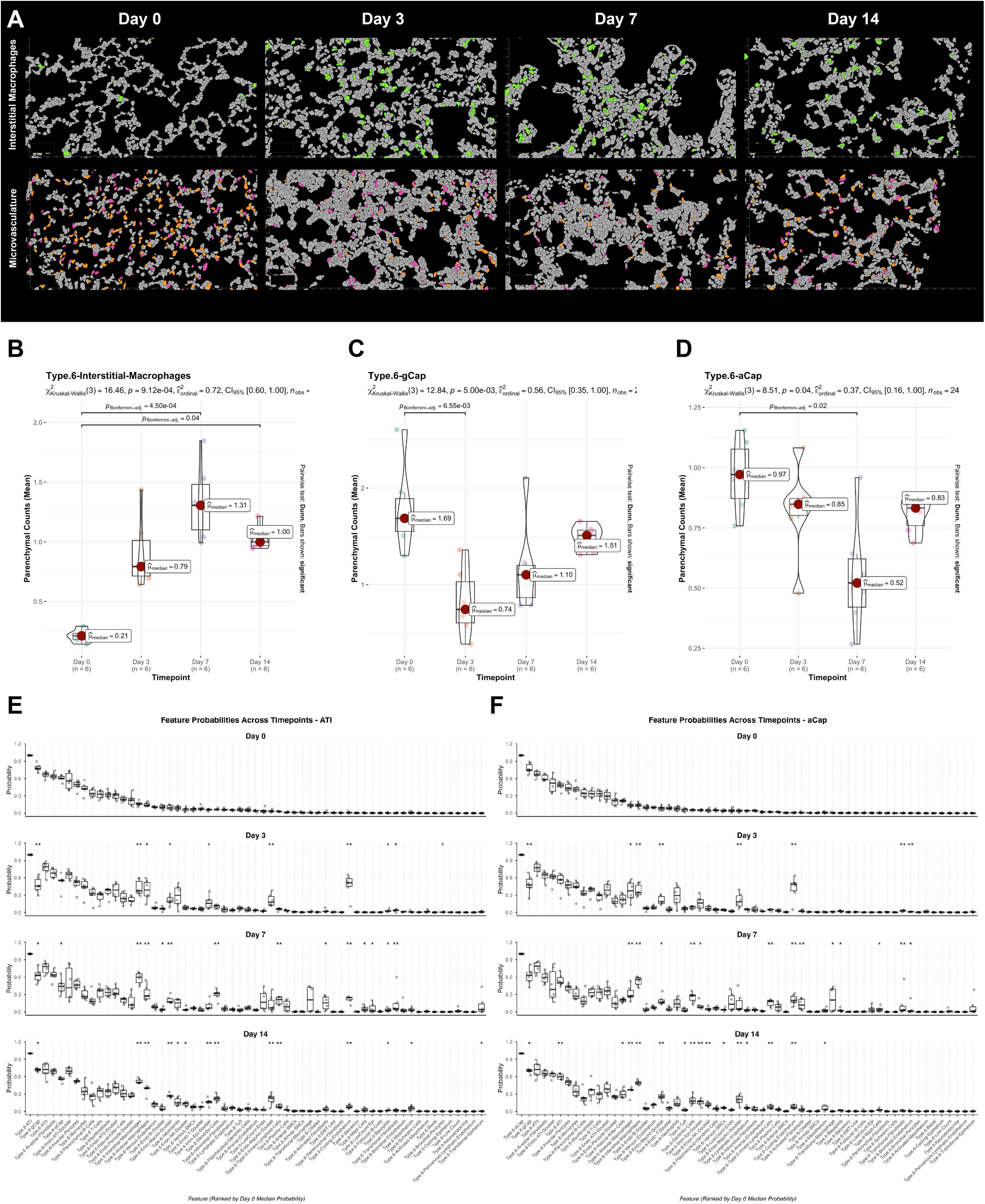
Immune dynamics over time. **(A)** Representative regions from Day 0, 3, 7, and 14, with interstitial macrophages (top, green) and the aCap and gCap capillary subtypes (bottom, highlighted) shown against the underlying tissue graph (grey). Interstitial macrophage density increases in the parenchyma by Day 3, remains elevated during Day 7 remodeling, and tapers by Day 14, a pattern spatiotemporally coupled to local loss of well-differentiated microvasculature (bottom row). **(B)** These trends are significant and conserved across replicates (*n* = 6 per timepoint). Interstitial macrophage counts rise from a Day 0 median of 0.21 to a Day 7 peak of 1.31 before declining to 1.00 by Day 14 (Kruskal-Wallis χ²(3) = 16.46, *p* = 9.12×10⁻⁴); Day 0 differs significantly from both Day 7 and Day 14 (Dunn’s post hoc, Bonferroni- adjusted *p* = 4.50×10⁻⁴ and 0.04, respectively). **(C)** gCap density falls significantly from Day 0 to Day 3 (median 1.69 to 0.74; χ²(3) = 12.84, *p* = 5.00×10⁻³; Bonferroni-adjusted *p* = 6.55×10⁻³), then partially recovers by Day 7 and Day 14 (medians 1.10 and 1.51). **(D)** aCap density declines progressively to a Day 7 nadir (median 0.52, down from 0.97 at Day 0; χ²(3) = 8.51, *p* = 0.04; Bonferroni-adjusted *p* = 0.02), then partially recovers by Day 14 (median 0.83). **(E,F)** Neighborhood composition analysis, based on edge colocalization, for ATI (E) and aCap (F) cells, with candidate neighbor features ranked by Day 0 median probability. Relative to the Day 0 benchmark, Day 3, 7, and 14 show significant compositional shifts in both neighborhoods (asterisks, pairwise tests vs. Day 0), marked by a loss of microvascular neighbors and enrichment of myeloid and lymphoid neighbors, consistent with the immune infiltration and capillary rarefaction quantified in (A-D). Notably, this coupled macrophage expansion and capillary loss is transient, beginning to resolve by Day 14 rather than persisting, consistent with this model’s broader distinction from IPF, in which an analogous fibrogenic-like community fails to resolve.

**Figure S11.**
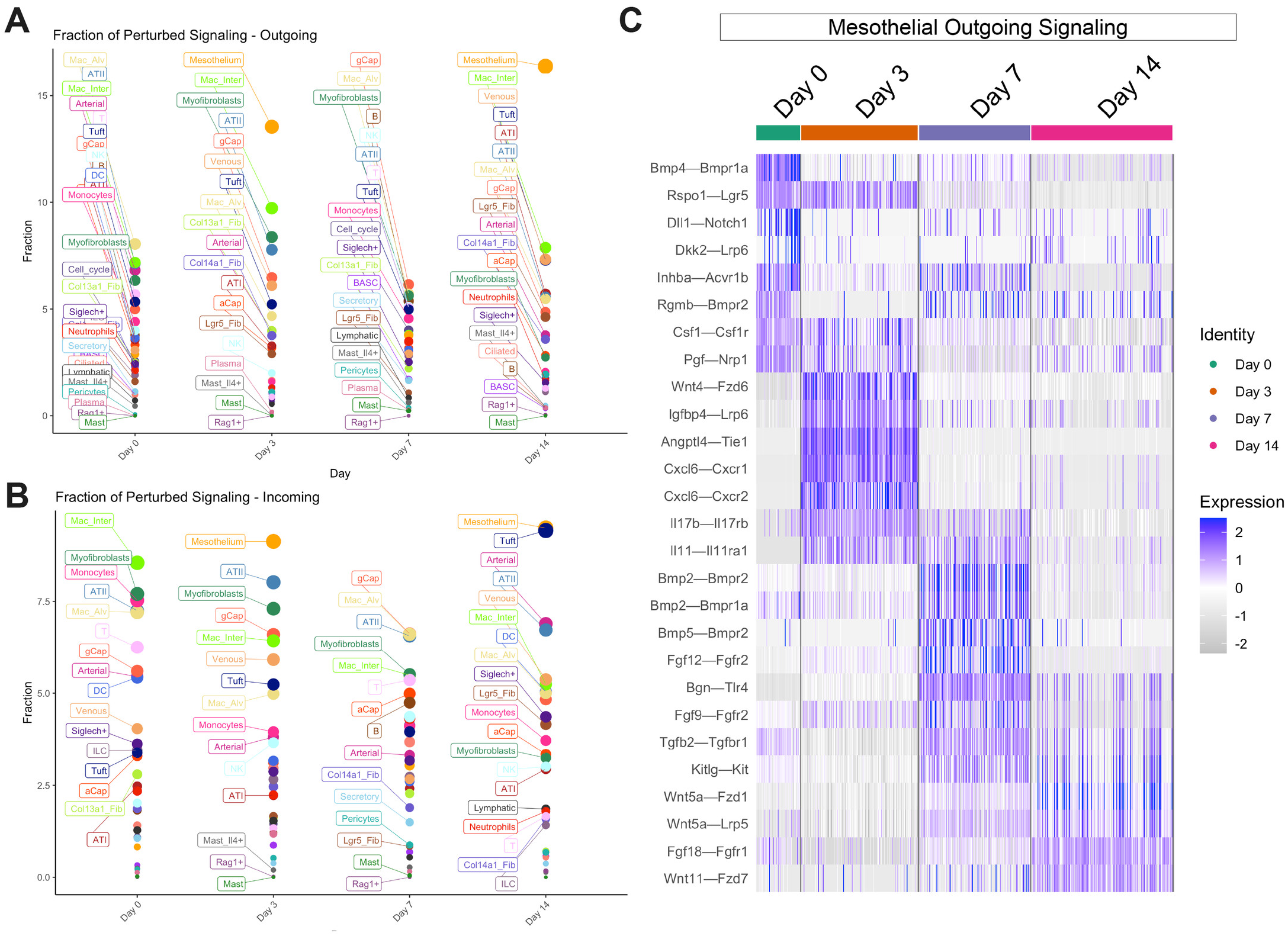
Mesothelial interactome over time. **(A,B)** Fraction of perturbed signaling, ranked by cell type, for outgoing (A) and incoming (B) signaling at each timepoint. The mesothelium is the top-ranked perturbed node for both outgoing and incoming signaling at Day 3 and Day 14, across all cell types profiled. **(C)** Heatmap of mesothelial outgoing ligand-receptor signaling, at single-cell resolution, grouped by timepoint. Distinct temporal blocks are apparent: developmental/morphogen pathways (e.g., *Bmp4*-*Bmpr1a*, *Rspo1*-*Lgr5*, *Dll1*- *Notch1*, *Dkk2*-*Lrp6*) are most active at baseline (Day 0); *Wnt4*-*Fzd6*, *Igfbp4*-*Lrp6*, *Angptl4*-*Tie1*, and both *Cxcl6* receptor pairs (*Cxcr1*, *Cxcr2*) form a cluster restricted to Day 3; *Bgn*-*Tlr4* and *Fgf9*-*Fgfr2* mark a second cluster enriched at Day 7; additional shifts emerge by Day 14. The Day-3-restricted ligands (*Wnt4*, *Igfbp4*, *Angptl4*, and *Cxcl6*) are established regulators of tissue remodeling, and several are also known regulators of normal lung development, positioning the mesothelium as a candidate early orchestrator of the regenerative response; *Bgn* and *Fgf9* extend a similar remodeling-associated signature into the Day 7 fibromimetic window.

**Figure S12.**
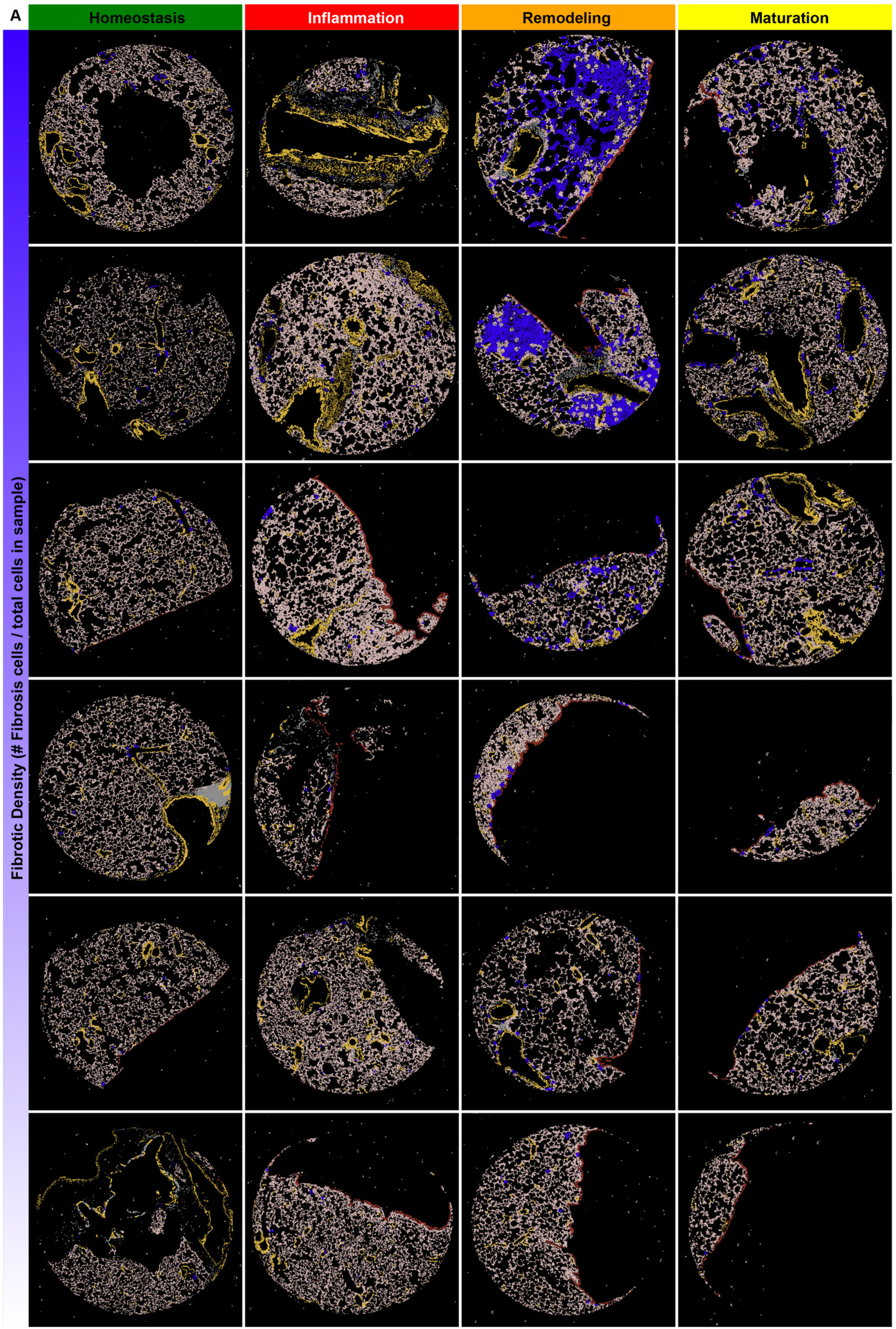
Fibrosis density dynamics across the PPLR time course. (A) Cell-resolved spatial maps of representative tissue pucks at four phases of post- pneumonectomy lung regeneration — Homeostasis (Day 0), Inflammation (Day 3), Remodeling (Day 7), and Maturation (Day 14). Each puck shows individual segmented cells colored by category (pink, parenchymal cells; yellow, conducting airway; purple, fibrotic / transitional remodeling community cells; red, pleural). Pucks are arrayed vertically by fibrosis density (defined as the fraction of fibrosis-category cells out of total cells per puck), with highest-density samples at top and lowest at bottom. The Remodeling phase (Day 7) shows the highest fibrosis density and is the only phase in which substantial spatial patches of fibrosis cells are observed; by Maturation (Day 14), fibrosis density returns to near-homeostatic levels. Fibrosis density varies between pucks within a given phase, reflecting the spatial heterogeneity of the transitional remodeling community.

**Figure S13.**
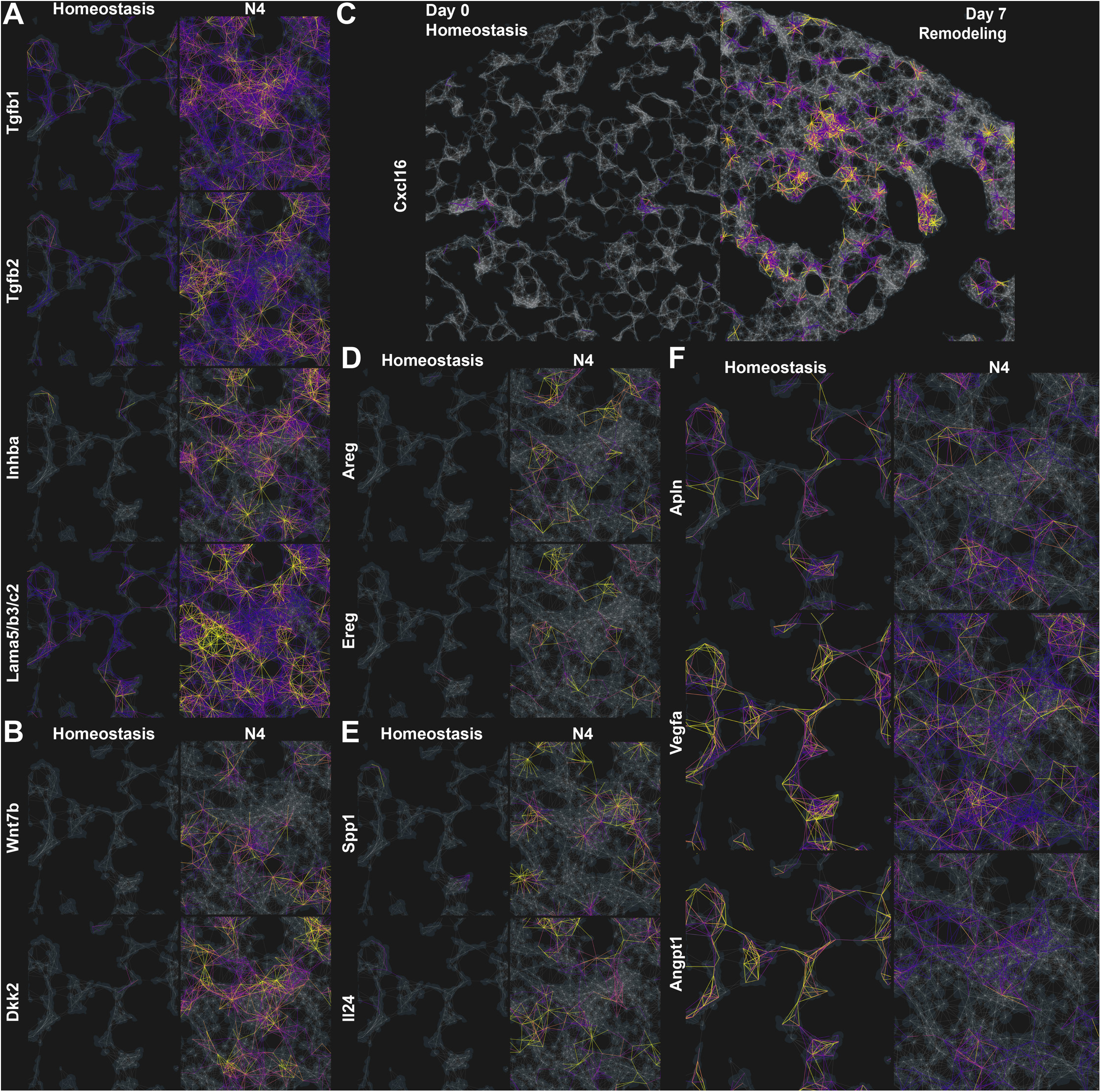
N4 connectivity. Selected single-feature examples from the N4 fibromimetic connectivity cluster, contrasting homeostatic tissue with N4 regions (Day 0 vs. Day 7 in C). **(A)** N4 regions show marked upregulation of *Tgfb1* and *Tgfb2* connectivity, along with *Inhba*-mediated connectivity and connectivity mediated by the laminin chains *Lama5*, *Lamb3*, and *Lamc2*. **(B)** Wnt family signaling is also strongly perturbed, exemplified by the activator *Wnt7b* and the inhibitor *Dkk2*. **(C)** These regions sit within a broader tissue context marked by prominent inflammation, including non- homeostatic *Cxcl16* connectivity across the Day 7 remodeling lung relative to Day 0. **(D)** Actively remodeling regions show significant upregulation of amphiregulin (*Areg*) and epiregulin (*Ereg*) connectivity. **(E)** These regions also show upregulation of *Spp1*- and *Il24*-mediated connectivity. **(F)** In contrast, active remodeling is associated with a loss of connectivity linked to maintenance of well-differentiated microvasculature, shown here for the *Apln*, *Vegfa*, and *Angpt1* families.

